# Structural and functional characterization of a putative *de novo* gene in *Drosophila*

**DOI:** 10.1101/2021.01.18.427054

**Authors:** Andreas Lange, Prajal H. Patel, Brennen Heames, Adam M. Damry, Thorsten Saenger, Colin J. Jackson, Geoffrey D. Findlay, Erich Bornberg-Bauer

## Abstract

Comparative genomic studies have repeatedly shown that new protein-coding genes can emerge *de novo* from non-coding DNA. Still unknown is how and when the structures of encoded *de novo* proteins emerge and evolve. Combining biochemical, genetic and evolutionary analyses, we elucidate the function and structure of *goddard*, a gene which appears to have evolved *de novo* at least 50 million years ago within the *Drosophila* genus.

Previous studies found that *goddard* is required for male fertility. Here, we show that Goddard protein localizes to elongating sperm axonemes and that in its absence, elongated spermatids fail to undergo individualization. Combining modelling, NMR and CD data, we show that Goddard protein contains a large central α-helix, but is otherwise partially disordered. We find similar results for Goddard’s orthologs from divergent fly species and their reconstructed ancestral sequences. Accordingly, Goddard’s structure appears to have been maintained with only minor changes over millions of years.

## Introduction

*De novo* evolved genes are novel genes that arise from previously non-coding DNA [1, 2, 3, 4]. Contrary to most other newly evolved genes, which are generated by duplications [5] or rearrangements of existing gene fragments [6], *de novo* genes are not derived from existing, protein-coding sequences. Accordingly, selection may only act on the functional structure of an encoded protein after it has been born. *De novo* genes have been confirmed across a wide range of eukaryotes [7, 8, 9, 10, 11, 12, 13, 14].

Studies over the last decade have illustrated the key mechanisms underlying these genes’ emergence. Many *de novo* protein-coding genes start out as a non-coding transcripts, transcribed from intergenic or intronic regions. As circumstantial events, mutations that create a protein-coding open-reading frame (ORF) can occur in the transcribed region [15, 16, 11, 17, 13, 14]. The order of these steps can vary, and each of the steps is frequent.Recent evidence showed an abundance of species-specific, newly generated transcripts supporting *de novo* gene emergence [18, 19, 11]. While most novel transcripts are quickly lost, those that are retained and encode a polypeptide are exposed to selection, eliminating novel proteins that are deleterious for cell function [20, 11]. While the computationally predicted structural properties (such as secondary structure and disorder) of *de novo* proteins do not appear to change significantly over millions of years [11], relatively little is known about how these properties are acquired upon gene birth. In particular, it remains unclear how *de novo* genes, which arise from essentially random sequences, are able to form their initial structures, acquire functions, become fixed in a population, and persist beyond several speciation events. Understanding these issues is important, given that *de novo* genes have already changed our perceptions of how genomic novelties can arise [21]. In particular, it is often proposed that newly evolved genes in general, and *de novo* genes in particular, are involved in many important processes such as development, stress response and environmental adaptation [22, 4]. Most insights concerning *de novo* gene evolution stem from large scale comparative genomic, transcriptomic and proteomic studies. Less is known about the specific functions and structures of *de novo* proteins, because very few of them have been studied in detail. One example is the ‘antifreeze glycoprotein’ (AFGP), which protects Arctic codfishes from freezing [23, 24]. AFGP acquired, probably by convergent evolution, a structure that is similar to an evolutionary unrelated antifreeze protein found in Antarctic notothenoids [25, 26]. Another example is Bsc4, a non-essential *de novo* protein found in *Saccharomyces cerevisiae* and implicated in DNA repair under nutrient-deficient conditions [8, 27]. Bsc4 contains large disordered regions [28], but further details regarding its structure, cellular location and function remain unclear. Recently, a *de novo* yeast protein was computationally and experimentally shown to progressively evolve properties that place it into the endoplasmic reticulum membrane [29]. Finally, two putative *de novo* genes, named *goddard* (*gdrd*) and *saturn*, have been identified in fruit fly [30] (note that we stick to a widely used convention in fly genetics, capitalising protein and putting the gene names in lower case italics). Both genes appear to have arisen from intronic regions at least 50 million years ago (Mya), at the root of the *Drosophila* genus (**Figure S1**), and preliminary structural features have been predicted computationally. Both genes are expressed specifically in the male reproductive tract, a pattern conserved across many species, and RNA interference tests found that both are essential for male fertility in *D. melanogaster* (*Dmel*)[30].

A recent conceptualisation for defining *de novo* gene functionalisation [31] describes five levels of functional analysis. Our previous work on *gdrd* [30] addressed the gene’s expression (a conserved, male-specific pattern across *Drosophila* species) and began to investigate the protein’s evolutionary implications (conserved in most fly species except for *D. willistoni*) and capacities predicted to have one major α-helix with disordered termini). Here, we present a more detailed analysis of the structure, function, and evolution of the Gdrd protein, which allowed us to describe its molecular and structural properties, its cellular function and thus its potential, physiological implications and further elaborate on its evolutionary implications [31]. We first use computational and experimental approaches to determine the structure of Gdrd protein from *Dmel*. Then, using null and tagged rescue alleles of the *gdrd* gene, we show that Gdrd protein localizes to elongating sperm axonemes and it is required to form individual sperm cells in the post-meiotic testis. Finally, we predict the structures of orthologous Gdrd proteins from other *Drosophila* species and use ancestral sequence reconstruction to infer how this structure might have arisen and subsequently evolved. Gdrd’s high degree of structural conservation, coupled with its functional role, suggests that it likely became involved in spermatogenesis early in the evolution of the *Drosophila* genus.

## Results

### Gdrd is monomeric, soluble and compact with a helix at its core and disordered termini

To further assess the likelihood that Gdrd (with length 113 residues) forms a stably folded protein, we carried out predictions for a number of biophysical properties (see Materials and Methods). The *Dmel* Gdrd has a theoretical pI of 4.25 and does not contain any cysteine or tryptophan residues. Secondary structure predictions consistently indicate an α-helix at the core (residues 40-80), as suggested by Gubala et al. [30] (**Figure S2a-c**). Kyte-Doolittle hydrophobicity indicates 17% hydrophobic residues, which are primarily present in the core α-helix, with no indication of transmembrane regions (**Figure S2d**). Therefore, Gdrd is likely a stably folded protein.The existence of this core α-helix is further corroborated by the prediction of a coiled coil formation (CC) between residues 45 and 80. Consistent with the gene’s potential *de novo* origin, CCs can be formed relatively easily, from sequences which are almost random, provided they have at least a clear hydrophobic-polar pattern [33]. Generally, many predicted CCs, in particular short ones, have overlapping predictions, e.g. with regions predicted to be disordered [33]. However, this ambivalence may also reflect their true structural state, since CCs are often formed from non-folding structural elements in response to triggers such as binding to another protein [33, 34]. A second, shorter helix is also predicted near the N-terminus (residues 10-18), but with lower confidence. The rest of the protein, in particular the termini, is predicted as disordered [30]. Using additional programs (see Material and Methods) to investigate Gdrd even further, we find: (i) that the core α-helix is stably folded, while the termini of the protein appear less ordered, (ii) no indication of toxic aggregation-prone segments [4], (iii) that solubility is predicted to be high for much of the sequence with the exception of the hydropathic core helix, (iv) finally, that Gdrd is not predicted to have regions likely to undergo liquid-liquid phase separation. Taken together, preliminary structural predictions indicate that Gdrd adopts a soluble (validated by SDS-PAGE, **Figure S2f**), relatively compact, non-aggregating structure with helices at its core (**Fig S2a-d**) and N-terminus and partially disordered regions throughout the remainder of the protein(**Figure S2 and S3**).

We further corroborated these predictions with *ab initio* tertiary structure prediction using the QUARK server [35]. Consistent with the above heuristic methods, a helical core and terminal disorder are predicted (**Figure S4**). We performed a pairwise root-mean-square deviation (RMSD) analysis between the top predicted Gdrd structure and the following four top structures and found that the major predicted structural features of Gdrd are conserved across all five top structures. While these all-atom RMSD values are relatively high, ranging from 11 to 13 Å, such values are reasonable considering the flexible Gdrd C-terminus. Additionally, we observe a considerably lower RMSD for the core helix, for which all pairwise RMSD measurements are < 3 Å (**Figure S4**) [36]. We also applied the previous structural prediction methodology to a 6x-His-tagged version of Gdrd from *Dmel* and obtained highly similar results to the untagged Gdrd protein, a finding that supports the use of tagged Gdrd for further experimental work (**Figure S5**). Using the top predicted structure from QUARK as an input template (**Figure S3b**), we then performed three independent MD simulations using Gromacs [37, 38, 39] (see Materials and Methods). Across all three simulations the structures rapidly diverge from the input template and reach relatively high RMSD values (Figure S3) due to significant disorder in the termini and loop regions. However, the central helix and a portion of the N-terminus remain stably folded (**Figure 1a**). A residue-by-residue RMSF analysis confirms these results, demonstrating greater rigidity in the central helix and N-terminus than throughout the rest of the protein across all three simulations (**Figure 1b and c**).

**Figure 1:**
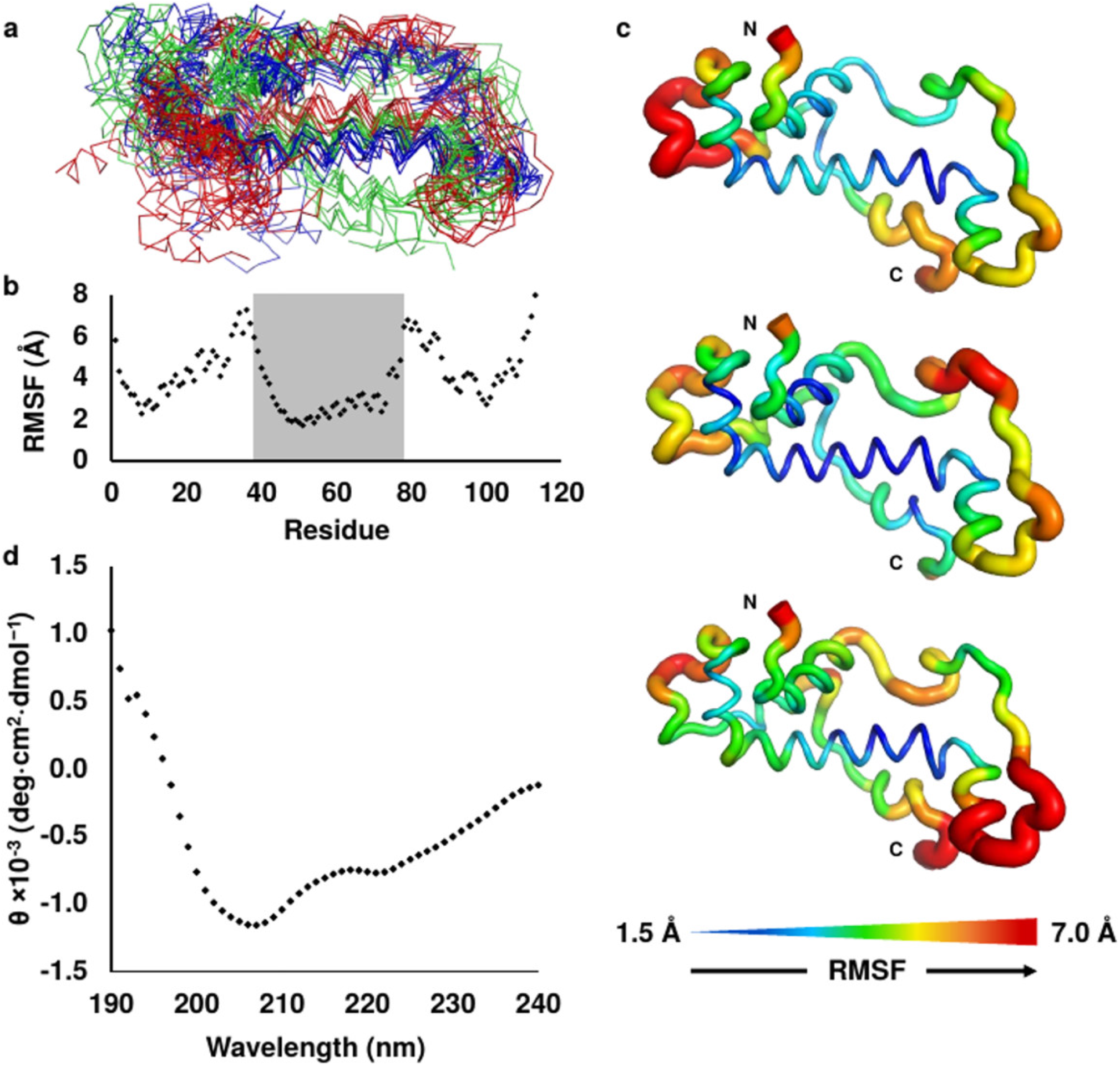
Molecular Dynamics (MD) and Circular Dichroism (CD) of Gdrd confirm a partially ordered helical structure. a) Representative backbone ensemble of the modeled Gdrd structure composed of ten frames sampled every 20 ns from each of three 200 ns MD replicates (shown as green, blue, and red ribbons respectively). The central helix and a portion of the N-terminal helix remain stably folded across all three simulations despite considerable flexibility in the rest of the protein structure, indicative of a partially ordered structure. b) Plot of C_*α*_ Root Mean Square Fluctuation (RMSF) versus residue position (averaged over three MD replicates) further demonstrates that the central helix of Gdrd, shaded, is the most conformationally rigid structure in the protein (C_*α*_ RMSF = 2-4 Å). c) Mapping the RMSF values to a representative Gdrd MD structure for each of the three simulations shows similar regions of conformational flexibility for each replicate. d) CD spectrum of Gdrd demonstrates characteristics typical of helical proteins. A helix minimum at 222 nm that is weaker than the helix minimum at 208 nm is characteristic of a flexible or distorted helix as was observed in the MD simulation [32].

In order to assess the novelty of the structures predicted for Gdrd, we used 3D-BLAST and mTM-align to compare our *ab initio* models of Gdrd to all structures in the Protein Data Bank (PDB) [40, 41, 42]. With both methods, we find no clear similarity to any known eukaryotic structures when searching the top five models predicted for Gdrd against the PDB − but note that Gdrd’s short length results in a large number of spurious alignments of Gdrd’s helix-turn motif with the helical bundles of larger, unrelated proteins.

To confirm our computational predictions, we cloned and overexpressed Gdrd in *Escherichia coli* (strain BL21 Star(DE3)). We note that expression attempts using a range of tags (Maltose Binding Protein, Strep-tag and the Fh8 tag) and restriction sites were unsuccessful, reminiscent of the complications encountered in expressing Bsc4 [28]. For Gdrd, some fractions failed to elute, while others formed inclusion bodies which could not be further purified (see Materials and Methods). Only the combination of an N-terminal 6x-His tag with purification over a Nickel column yielded exploitable concentrations of >8 mg/mL soluble protein (**see Figure S2f**). Mass spectrometry (see online supplementary data, Zenodo DOI: 10.5281/zenodo.4291827) indicate that Gdrd is monomeric in solution. Accordingly, if the helical core region does indeed form a CC, the CC does not mediate homodimerization [34]. However, this does not rule out a role for the CC in heterodimerization with other proteins or binding to small molecules. Experimental far UV circular dichroism (CD) spectra of purified Gdrd comply with our computational predictions: the two minima at 222 nm and 208 nm indicate a flexible α-helical character, while the absence of a minimum around 218 nm suggests a lack of *β*-sheet regions (**Figure 1d**). CD results also confirm that Gdrd is at most partially disordered, since highly disordered proteins are expected to show negative ellipticity below 210 nm, with a minimum at 195 nm [43, 44]. For Gdrd, the CD signal shows positive ellipticity below 200 nm. Indeed, computational interpretation of the CD spectrum using K2D3 suggests an α-helical content of 85% [45], further supporting the non-transient nature of a shorter N-terminal helix in addition to the longer helix at the core of Gdrd. Also consistent with both computational predictions (Quark, MD) and CD spectroscopy, the ^15^N-HMQC NMR spectrum indicates that a large region of Gdrd is partially disordered, as indicated by broadened or missing peaks, with the remainder of the protein showing a spectrum characteristic of a low-diversity fold, as would be expected for a primarily helical rigid region (**Figure S6**). Last, we performed a thermal unfolding experiment (thermal shift assay, TSA) using SYPRO Orange (**Figure S7**). Consistent with CD and NMR, the TSA of Gdrd shows a thermal unfolding transition at 47.3°C +/-0.9°C, indicating that Gdrd possesses a fold that can be denatured, exposing additional dye binding sites. Characterization by fluorescence and re-folding experiments were not performed due to the lack of tryptophan or cysteine residues, and *β*-sheet content.

### Gdrd is required for male fertility

Having examined the structure of Gdrd, we next attempted to clarify its ‘physiological implications’ and ‘interactions’ of *gdrd* [31]. Initial phenotypic characterization of *gdrd* using testis-specific knockdown revealed that *gdrd* was required for the production of mature sperm [30]. This initial finding suggested that *gdrd* functions during spermatogenesis. To better understand the functional role of *gdrd* in fertility and bypass any potential drawbacks of RNAi, such as partial gene knockdown, we generated null alleles using CRISPR-Cas9 genome editing [46]. Using two guide RNAs to simultaneously target loci 627 bp upstream of the *gdrd* start site and 278 bp downstream of the stop codon, we generated two independent alleles, both of which completely deletes a Gdrd protein-coding region (**Figure 2a**).

**Figure 2:**
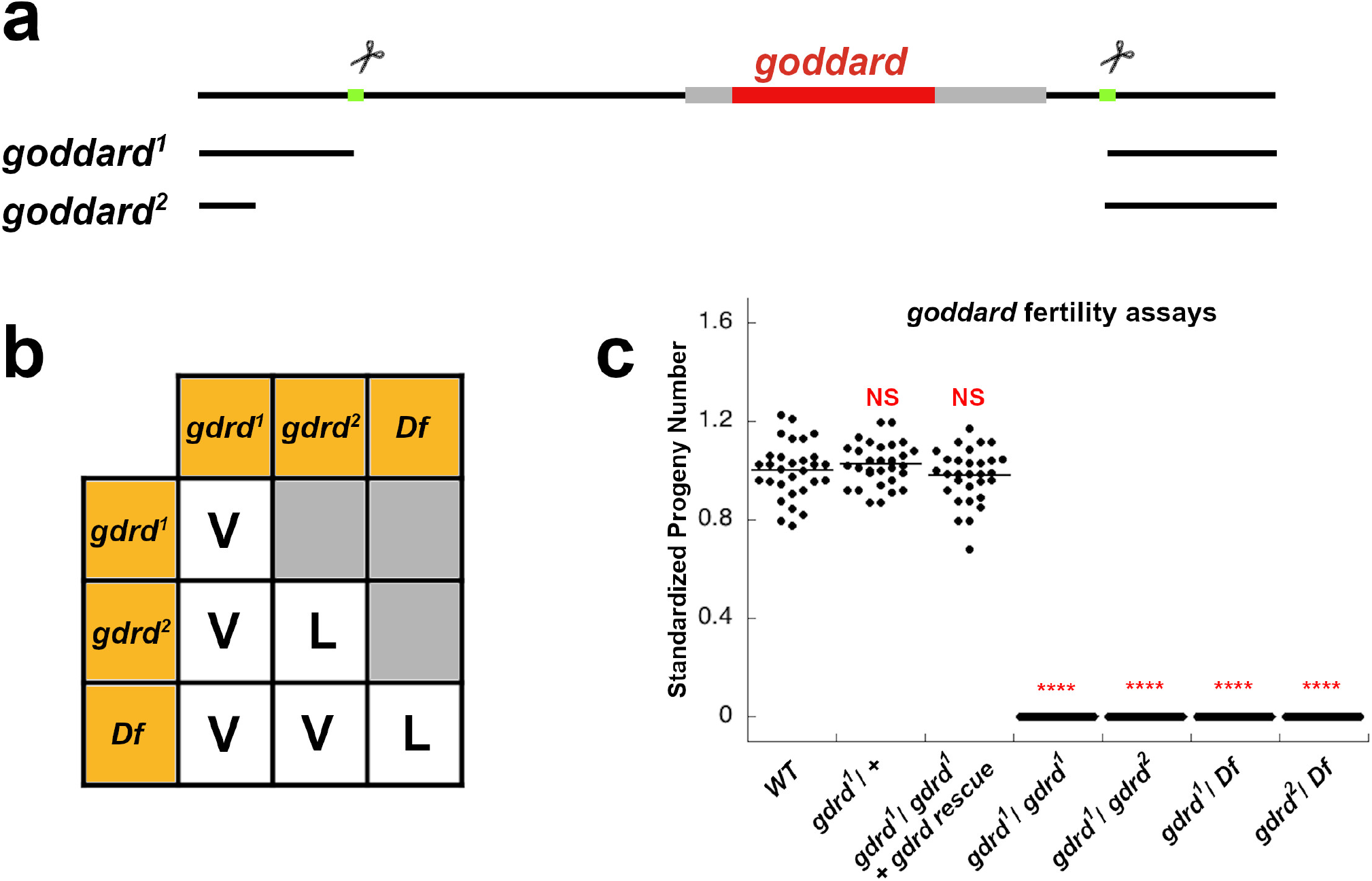
*gdrd* null alleles are viable but cause male sterility. (a) *gdrd* genomic locus. Two CRISPR-Cas9 target sites (green) that flank the *gdrd* CDS (red) were selected to create null alleles of the gene. The *gdrd*^*1*^ allele results from a precise deletion at both sites, whereas the *gdrd*^*2*^ allele deletes an additional 153 bp upstream of the 5’ target site. (b) Complementation test using the *gdrd* alleles and a large, molecularly defined deficiency *Df(3L)ED4543* that spans the *gdrd* locus shows that loss of *gdrd* has no effect on organismal viability; V= viable and L= lethal. (c) *gdrd* mutants as homozygotes or in heterozygous combinations with each other or a deficiency *Df(3L)ED4543* are all 100% sterile (n=30 for all genotypes). Standardized average progeny number (progeny number/ average progeny number) was 1.00 ± .110 (± se) for WT flies (*w*^*1118*^), 1.03 ± .087 (± se) for *gdrd* /+ heterozygous flies, and .98 ± .111 (± se) for *gdrd* rescue flies (homozygous *gdrd* mutants carrying a single copy of the *gdrd* rescue construct). For all *gdrd* loss of function backgrounds, average progeny number was 0 ± 0 (± se). Statistical analysis: two sample T test, **** P<0.0001.

The *gdrd*^*1*^ allele deletes a 1.27 kb segment of the genome, reflecting a deletion generated by precise double-stranded cuts at the targeted sites (**Figure 2a**). A second allele, *gdrd*^*2*^, removes an extra 153 bp upstream of the 5’ target site, forming an even larger deletion (**Figure 2a**). We found that *gdrd*^*1*^ homozygous mutant flies are viable, while *gdrd*^*2*^ homozygous mutant flies are lethal (**Figure 2b**). Flies carrying either mutant allele in combination with a molecularly defined deficiency that spans the *gdrd* gene or in combination with each other are, however, viable, suggesting that the chromosome bearing the *gdrd*^*2*^ mutation also has a second site lethal mutation not associated with the *gdrd* gene.We further confirmed that *gdrd*^*1*^ specifically affects the *gdrd* locus by analyzing the mRNA levels of both *gdrd* and a neighboring locus, *CG5048* (**Figure S8**). Altogether, these data suggest that *gdrd* is not required for organismal viability, consistent with the observation that the gene is primarily expressed in the testis.

We next used various combinations of *gdrd* mutant alleles and the defined deficiency line to replicate the results of the previous RNAi-based fertility assay [30]. The previous assay showed that depletion of *gdrd* transcripts within the male germ cell lineage causes complete sterility. Likewise, our *gdrd* alleles, either as homozygotes or in combination with each other or a deficiency, are also fully sterile, suggesting that our mutations are functionally null alleles (**Figure 2c**).

### Gdrd is expressed in elongating spermatid cysts and associates with growing axonemes

To investigate the expression pattern and subcellular localization of the Gdrd protein *in vivo*, we generated a hemagglutinin (HA)-tagged *gdrd* genomic rescue construct containing both the Gdrd coding region and its upstream and downstream regulatory elements. This construct, when placed into a *gdrd* mutant background, is sufficient to restore fertility to wild-type levels, indicating that the construct produces functional Gdrd proteins at sufficient levels and in the correct spatio-temporal pattern during spermatogenesis (**Figure 2c**). Using an antibody (AB) that recognizes the HA epitope, we then visualized HA-tagged Gdrd proteins in whole mount testes. Gdrd expression most likely starts in mature spermatocytes and peaks in early spermatids (**Figure 3a-a”**). Interestingly, Gdrd expression isn’t observed at the basal end of the testis, where mature individualized sperm are present, indicating that Gdrd is transiently expressed primarily during sperm tail elongation stages of spermatogenesis (**Figure 3a-a”**).

**Figure 3:**
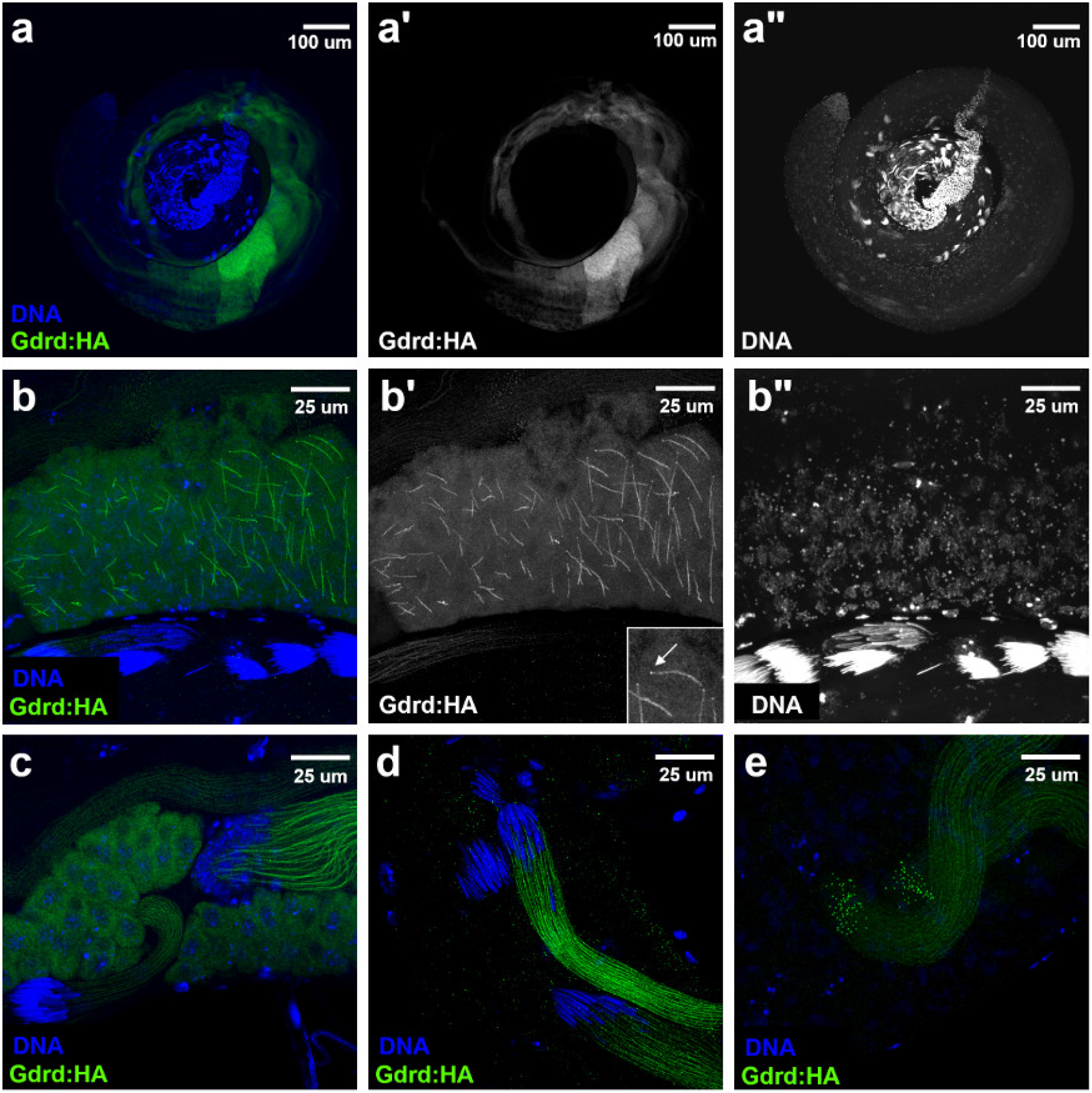
Gdrd protein expresses during spermatid elongation and localizes to the growing axoneme. (a-a”) Gdrd protein expression turns on during spermatid elongation. HA-tagged Gdrd protein expressed from a functional rescue construct in a *gdrd* mutant background is present weakly in mature spermatocytes and peaks in the early spermatids. Gdrd expression, however, is not present at the basal end of the testes, indicating that the protein is not present in individualizing or mature spermatids. (b-b”) Gdrd localizes to the growing axoneme during early stages of spermatid elongation. Inset in (b’) shows that Gdrd localizes to the axoneme as well as a distally located punctate structure (arrow) that is reminiscent of proteins that localize to the ring centriole. (c) Gdrd localization in spermatid bundles with round nuclei (upper right) or canoe shaped nuclei (lower left) becomes exclusive to the axonemes. (d) Spermatid bundles at the basal end of the testes with canoe shaped nuclei express Gdrd whereas later staged spermatid bundles (with needle shaped nuclei) lose Gdrd expression. (e) Distal ends of two elongating spermatid bundles. Gdrd localizes to both the growing axonemes and distally located punctate structures.

An analysis of the protein’s subcellular localization indicates that Gdrd is primarily cytoplasmic. In mature spermatocytes, the protein shows a nuclear exclusion pattern, though a low level of protein may exist in the nucleus as well (**Figure 3a-a”**). Later during spermatogenesis, at the onset of sperm tail elongation, the localization of the Gdrd protein starts to shift from the cytosol to the growing axoneme (**Figure 3b-b”**). Indeed, in highly elongated spermatid bundles, Gdrd is undetectable in the cytosol and is exclusively associated with growing axonemes (**Figure 3c-d”**). During these later stages of elongation, the initially round spermatid nuclei undergo a morphological change, stretching first into a canoe shape and then finally into a thin, needle like structure [47]. Using these shape changes to developmentally stage spermatid cysts, we find that there is a gradual reduction in the intensity of Gdrd staining in spermatid cysts with canoe shaped nuclei when compared to less mature cysts with round nuclei (**Figure 3c-d”**). This may reflect either the active degradation of Gdrd or the titration of the protein as cyst size and axoneme length increase. We also find that elongating spermatid bundles at the needle stage no longer have Gdrd expression (**Figure 3c-d”**). Besides this axonemal localization, we also observe a punctate Gdrd positive structures at the distal end of each growing axoneme during both early and late spermatid elongation (**Figure 3b-b’, 3e**). The localization pattern is highly reminiscent of proteins present in the ring centriole/ transition zone, a site important for axoneme growth and remodeling [48, 49].

### Gdrd is required for sperm individualization

We next determined the stage at which Gdrd is required during spermatogenesis. We crossed into the *gdrd*^*1*^ background a transgene expressing Don Juan:GFP (Dj:GFP), a sperm-tail marker that is activated during the early stages of sperm tail elongation [50]. Dj:GFP-positive elongating/elongated sperm tails are present in both wild type and *gdrd* mutant testes, suggesting that post-meiotic spermatids are, at the very least, able to undergo sperm tail elongation (**Figure 4a-b**). We also find that sperm coiling at the basal end of the testis occurs normally in both wild type and *gdrd* null flies, indicating that the absence of sperm in the seminal vesicle is not due to a defect in sperm packaging (**Figure 4a-b**). Simultaneously, we tested whether loss of *gdrd* affects sperm individualization. Individualization complexes (ICs) assemble normally at the apical end of nuclear bundles in *gdrd* mutant testes, but they fail to travel down the length of the sperm, indicating a failure in sperm individualization (**Figure 4a-b**). This defect occurs in both 1-day old and 3-day old mutants (**Figure 4c**). Furthermore, *gdrd* mutant testes show a major reduction in the total number of ICs, suggesting either a delay in IC formation or that spermatogenesis has halted in the tissue due to failure of IC translocation in older cysts (**Figure 4d**). We next quantified the number of nuclear bundles associated with ICs. We find that 57% of wild type nuclear bundles are associated with investment cones while only 23% of nuclear bundles are undergoing individualization in *gdrd* mutant testes (z-test, p <.00001). Finally, we found that waste bags, extraneous cytoplasm culled during individualization, are completely absent in both 1- and 3-day old *gdrd* null testes, consistent with our observation that the individualization complexes fail to move down the length of the elongated spermatids (**Figure 4e**). Taken together, these data show that in the absence of *gdrd*, sperm individualization can sometimes initiate but consistently fails to complete.

**Figure 4:**
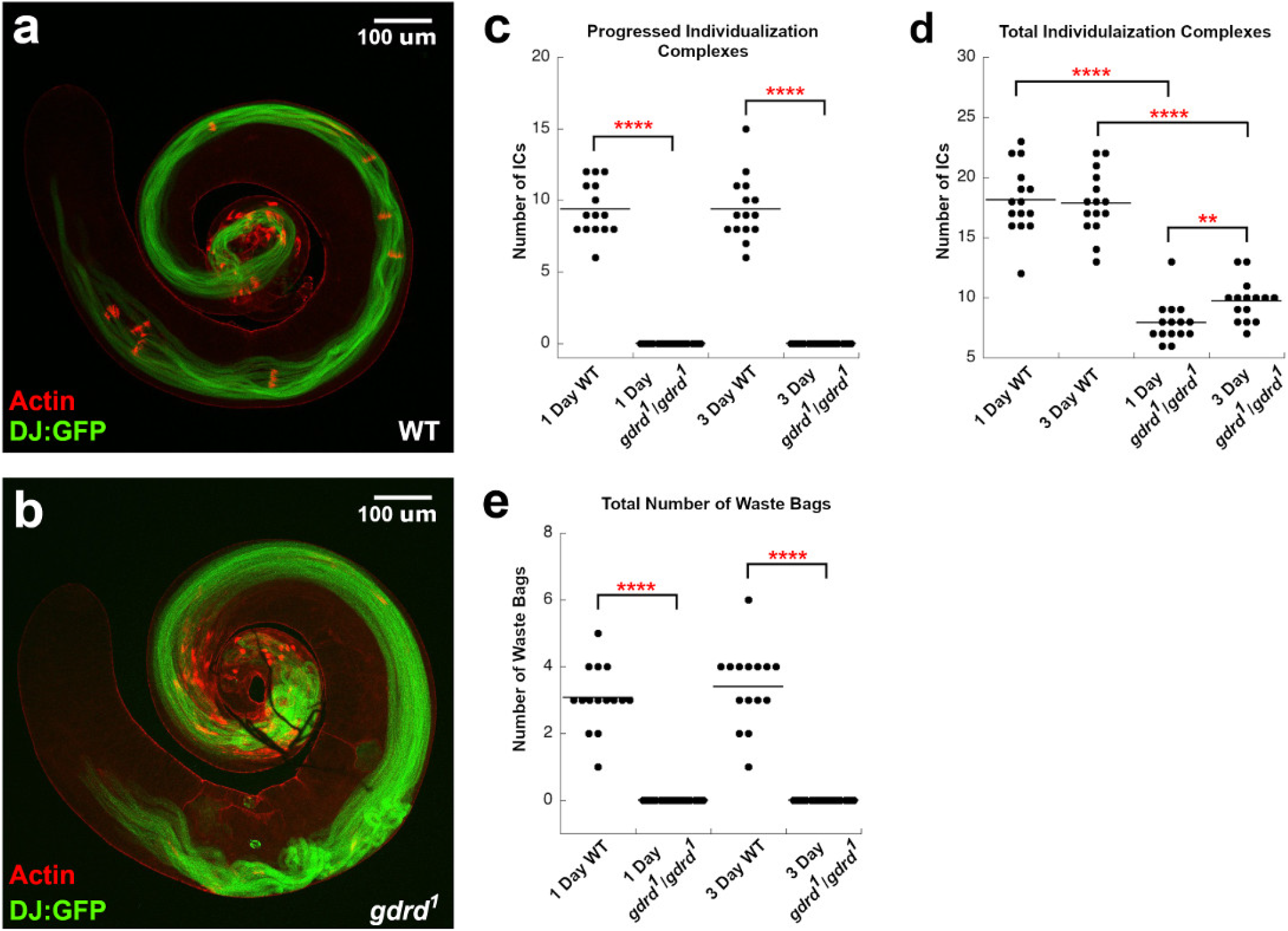
*Gdrd* mutant sperm fail to undergo individualization. (a) Wild type testis (*w*^*1118*^*/Y; dj:GFP/+*) and (b) *gdrd* mutant testis (*w*^*1118*^*/Y; dj:GFP/+*; *gdrd*^*1*^/*gdrd*^*1*^) were labeled with phalloidin to mark actin-rich individualization complexes (ICs). Dj:GFP (green) marks elongated/ elongating sperm. Scale bars in (a) and (b) = 100 um (c) ICs assemble but fail to progress in *gdrd* mutant testes. Analysis was conducted on Day 1 post-eclosion and on Day 3 post-eclosion. Average number of progressed ICs was 9.4 ± .48 (± se) and 9.4 ± .54 (± se) in WT day 1 and day 3 testes respectively (n=15). Average number of progressed ICs was 0 ± 0 (± se) and 0 ± 0 (± se) in *gdrd* day 1 and day 3 testes respectively (n=15). (d) Assembly of ICs is decreased in *gdrd* mutant testes. Average number of total ICs was 18.1 ± .74 (± se) and 17.9 ± .68 (± se) in WT day 1 and day 3 testes respectively (n=15). Average number of total ICs was 7.9 ± .44 (± se) and 9.7 ± .44 (± se) in *gdrd* day 1 and day 3 testes respectively (n=15). (e) Waste bags are absent in *gdrd* mutant testes. Average number of waste bags was 3.1 ± .25 (± se) and 3.4 ± .31 (± se) in WT day 1 and day 3 testes respectively (n=15). Average number of waste bags was 0 ± 0 (± se) and 0 ± 0 (± se) in *gdrd* day 1 and day 3 testes respectively (n=15). Statistical analysis: Mann-Whitney U test, ** P<0.01, **** P<0.0001

Another feature of late spermatogenesis is nuclear condensation, whereby histones are stripped from chromatin and replaced with protamines, thus allowing for the compaction of the paternal genome [51, 52]. In *Dmel*, nuclear condensation is associated with nuclear reshaping, a process that is coordinated with both sperm elongation and IC assembly [53]. We observe that protamination of nuclear DNA occurs at the basal end of the testis in both wild type and *gdrd* mutants (**Figure 5a-b**), and that mutant testes and controls showed no significant difference in the number of nuclear bundles marked with protamine-GFP (**Figure 5c**). Consistent with our findings above, we do, however, observe a decrease in the number of protamine-positive nuclear bundles associated with ICs (Mann-Whitney U test, p-value < .00001), further indicating that individualization is lost in *gdrd* mutant testes (**Figure 5d**). We also noted a significant (Mann-Whitney U test, p-value < .00001) decrease in fully condensed, protamine-GFP-positive sperm bundles at the far basal end of the testis, suggesting the possibility that some mutant sperm are targeted for destruction (**Figure 5e**). Consistent with this idea, we frequently observed protamine-GFP positive nuclear remnants at the very basal end (**Figure 5a-b**). This finding is consistent with the observation that there are no mature sperm in the seminal vesicle.

**Figure 5:**
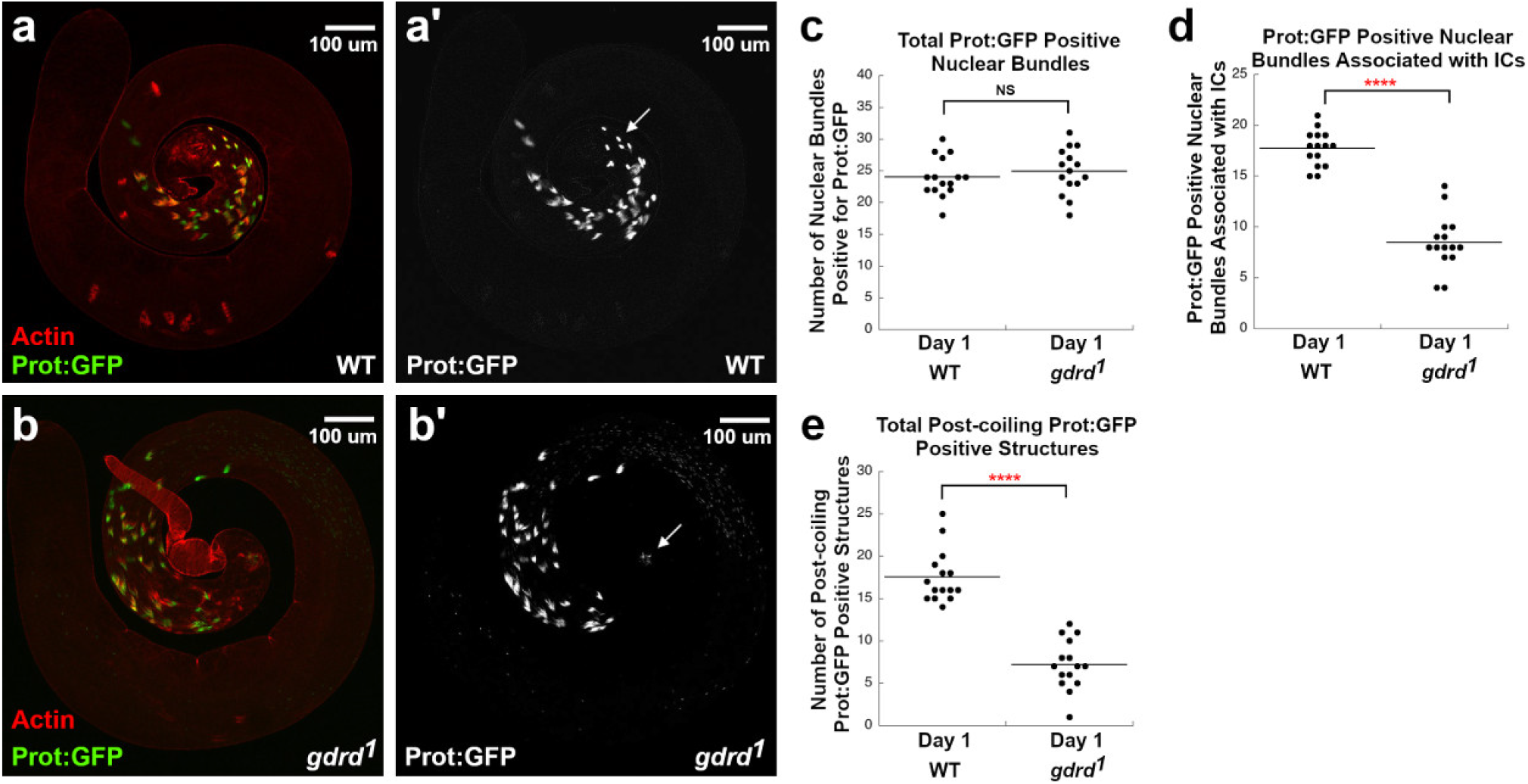
Nuclear compaction is normal in Gdrd mutant testis, but sperm bundles are potentially targeted for degradation post-coiling. (a) Wild type testis (*prot:GFP/Y; +/+*) and (b) *gdrd* mutant testis (*prot:GFP/Y* ;*gdrd* ^1^/*gdrd* ^1^) labeled with phalloidin (red) to mark actin-rich individualization complexes (ICs). Prot:GFP (green) labels protaminated nuclear bundles undergoing DNA compaction. (a’ and b’) Protamination of nuclei at the basal end *gdrd* mutant testis is unaffected. At the very basal end of nuclei, compact protamine positive structures associated with coiled sperm (arrows) are present in wild type (a’) testes but are mostly absent in *gdrd* mutant testes (b’). In *gdrd* mutant testis (b’), protamine positive remnants are present indicating that the bundled sperm may have undergone degradation. (c) Total number of protamine positive nuclear bundles is unaffected in *gdrd* mutants indicating that nuclear compaction is unaltered in *gdrd* mutants. Average number of protamine positive nuclear bundles was 24 ± .80 (± se) and 24.9 ± .93 (± se) in WT and *gdrd* day 1 testes respectively (n=15). (d) Association between protamine positive nuclear bundles and individualization complexes is decreased in *gdrd* mutant testis. Average number of protamine positive nuclear bundles associated with actin was 17.7 ± .45 (± se) and 8.5 ± .70 (± se) in WT and *gdrd* day 1 testes respectively (n=15). (e) The number of coiled sperm bundles is decreased in *gdrd* mutant testis, indicating that these structures are targeted for degradation. The n value for each sample was 15. Average number of post-coiling protamine positive structures was 17.5 ± .80 (± se) and 7.2 ± .76 (± se) in WT and *gdrd* day 1 testes respectively (n=15). Statistical analysis: Mann-Whitney U test, **** P<0.0001

### Structural properties of Gdrd have changed little since its birth and are conserved across the *Drosophila* genus

After looking into the possible structure of Gdrd and its function within *Dmel*, we next attempted to understand (i) if the properties of Gdrd have changed since its emergence and if so, (ii) how these changes might have influenced its function. Accordingly, we compared the structure of *Dmel* Gdrd to its orthologs from four *Drosophila* species: *D. ananassae* (*Dana*), *D. virilis* (*Dvir*), *D. mojavensis* (*Dmoj*), and *D. grimshawi* (*Dgri*) (**Figure 6 and S9**). We showed previously that Gdrd orthologs in the first three species listed are male-specific and testis-biased in expression, supporting a conserved role in male fertility [30]. Alignment of these sequences (**Figure 6 and S9**) demonstrates a conserved helical core between residues 40 and 79 of Gdrd, with average pairwise sequence similarity of 27.6% across the whole alignment and 51.6% within the core α-helix (excluding *Dgri*). To investigate the origins of this conserved core helix, we carried out ancestral sequence reconstruction (ASR) of Gdrd using orthologs from across the *Drosophila* clade (see Materials and Methods), and predicted the structures of the most likely ancestral sequences of (i) *Dmel*/*Dana*, (ii) *Dvir* /*Dmoj*/*Dgri*, and (iii) the common ancestor of all five species using Quark (**Figure 6**). Interestingly, the ancestor of *Dmel* and *Dana* is very similar to the two extant structures (115 aa, sequence similarity 78%). We repeated predictions of hydropathy, aggregation propensity, folding, and structural properties for all additional extant sequences and the three ancestors, with results almost identical to those of extant *Dmel* Gdrd (**Figure S1, S9, and S10**). Compared to Gdrd of *Dmel* and *Dana* the extended N- and C-terminal regions in *Dgri, Dmoj*, and *Dvir* are predicted to display shorter helices in addition to the core α-helix. In-terestingly, the ancestral central α-helix is predicted to be 10 aa shorter (28 vs. 38 aa) than the extant ones, suggesting that this helix has been gradually extended by accumulation of helix-stabilising mutations.

**Figure 6:**
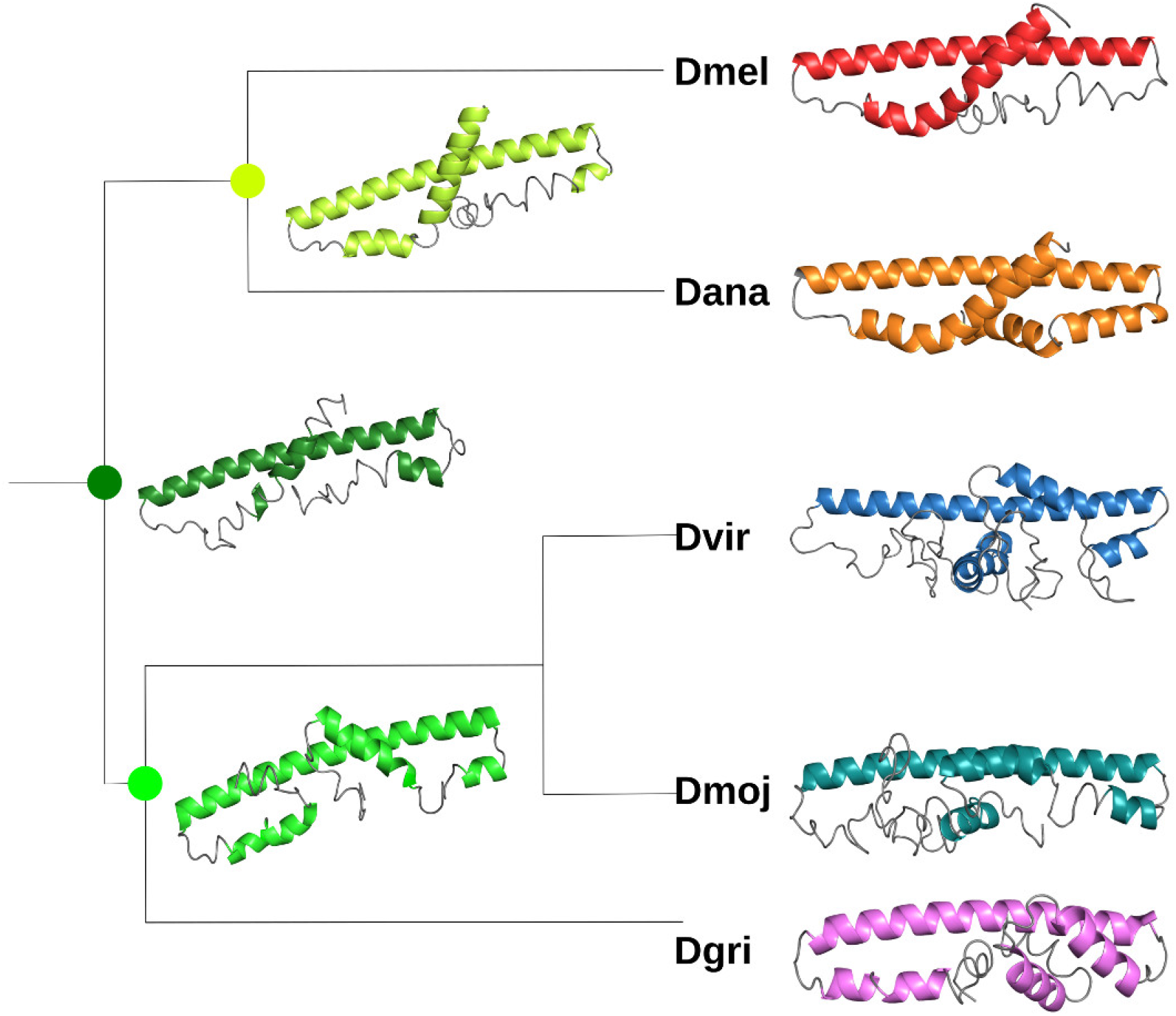
Structure prediction on ancestral reconstruction of *Gdrd* and its orthologs using Quark. Orthologs are from *Dmel, Dana, Dvir, Dmoj*, and *Dgri*. Additionally predictions for the most likely sequences for reconstructed ancestors of *Dmel*/*Dana* (bright green), *Dvir* /*Dmoj*/*Dgri* (green), and their most recent common ancestor (dark green) is shown. Helices are shown with a different colour in each species. PyMOL [54] was used to make protein cartoon structures (branch lengths are not meaningful).

Taken together, these results suggest that (i) the initial structure of Gdrd already featured a core α-helix upon gene emergence which (ii) was gradually extended during its early evolution but (iii) was largely unmodified over the last circa 15 million years, and (iv) terminal extensions forming short helices have been added in some but not all species.

## Discussion

Since their initial discovery more than a decade ago [7, 55, 8], *de novo* genes and the mechanisms underlying their emergence have been studied intensely. However, concerns regarding the reliability of their computational identification [56, 57, 58], and if and how they code for functional proteins, have also been raised. We showed previously [30] that Gdrd is likely a *de novo* evolved gene, supported by its absence from the syntenic regions of outgroup species, the lack of detectably similar proteins in any other taxa, its intronic location, and its high level of intrinsic disorder. Here, we combined multiple approaches, including compu-tational phylogenomic and structure predictions, experimental structural analyses, and cell biological assays, to further understand Gdrd’s structure, evolution, and importance in spermatogenesis.

In deducing the structure of Gdrd (based on *ab initio* protein structure prediction and MD, in combination with CD and NMR), we further confirmed the protein’s likely *de novo* origin, as structure-based homology searches using our models for Gdrd against all structures in the PDB detected no significant similarity in *Drosophila* and any other known eukaryotic species.We next observed that there is no transposable element (TE) nearby to the Gdrd locus (see Methods). A nearby TE remnant would indicate that *gdrd* could potentially be a strongly diverged transposed duplicate of another protein coding gene, which can also not be found in the genomes of outgroup species due to the disrupted synteny. Emergence from an intronic region thus remains the most plausible mechanism for the emergence Gdrd. How did the *gdrd* gene come to acquire its essential function in *D. melanogaster* ? We can gain clues from its structure and its generally high degree of evolutionary conservation within the genus. We found that the inferred ancestral form of the Gdrd protein has several intrinsically disordered (ID) regions, in addition to a helical folding core with a high predicted CC propensity. Rather than impeding protein functionality, ID is now recognised as an important structural feature that can mediate binding to a wide range of biomolecules and is occasionally essential for protein function [59]. Accordingly, the disordered termini of Gdrd may have helped the protein gain further interactions. Otherwise, Gdrd resembles what is believed to be a functional protein with rather average biochemical properties given that it has a folded core (supported by MD, CD, TSA, NMR), appears to be soluble, is not involved in phase separation (see Methods), and is neither aggregating nor multimeric. Terminal extensions of proteins with ID via loss and replacement of start or stop codons have been described before [60, 61] and may evolve into conserved stretches with domain-like properties over even longer time scales [62, 17]. Tretyachenko et al. [63] have also demonstrated that both ID and secondary structure can emerge from random sequences, much as *de novo* genes do. Likewise, ID may, to an extent, counter maladaptive aggregation [63]. Given that the protein properties of *de novo* proteins show a high degree of overlap with those of conserved and foldable proteins, it is therefore plausible that *de novo* proteins with properties such as (or similar to) Gdrd emerge from intergenic or intronic regions without prior adaptation. Based on its conserved structure (described here) and conserved male-biased expression pattern [30], we hypothesize that Gdrd was likely functional at or shortly after its emergence at the base of the *Drosophila* phylogeny, at least with respect to its expression and capacities. These levels have been described as the lowest level of function of *de novo* gene emergence [31].

The next levels of this model describe a new protein gaining interactions with other cellular components (e.g., proteins or membranes) and the acquisition of physiological implications for a specific biological process. Based on protein expression alone, *gdrd* most likely functions during the spermatid elongation phase of spermatogenesis. The protein’s expression starts in late spermatocytes and quickly peaks in early spermatids. While the initial localization of the Gdrd protein is predominantly cytosolic, the protein begins to associate with the growing axoneme at the start of spermatid elongation, suggesting a potential role for *gdrd* in regulating axonemal assembly. We also find that Gdrd expression is lost at the onset of individualization, indicating that Gdrd protein is not an integral component of the mature flagellum. Altogether, this expression pattern and localization suggest that Gdrd’s physical interactions are most likely restricted with proteins expressed during spermatid elongation. Our genetic analyses, which attempt to address the physiological role *gdrd* plays, indicate that the protein functions at or prior to spermatid individualization. Loss of *gdrd* leads to the arrest of spermatid cysts at the onset of individualization. While protamine expression and nuclear shape changes occur normally in *gdrd* mutant testes, fewer spermatid nuclear bundles are associated with ICs. This suggests that *gdrd* may be required to trigger individualization or its loss might affect some aspect of spermatid elongation itself such as axoneme growth, stability, and structure. Indeed genes that affect these processes often impair spermatid individualization [64, 49].

There are several cellular events that also correlate with the onset of individualization including IC assembly, activation of caspases, and the disassembly of the ring centrioles [65, 66, 67]. Interestingly, Gdrd localizes to a structure reminiscent of the ring centriole, a specialized area at the distal end of axonemes that coordinates axoneme growth and stability [67]. This localization thus raises the possibility that *gdrd* may function in regulating this structure. Another event that occurs during the transition from spermatid elongation to individualization is a switch from detyrosinated tubulin to polyglyclated tubulin [68, 49]. Gdrd likewise associates with axonemes during elongation and rapidly disappears at the onset of individualization, suggesting that Gdrd is most likely only associated with axonemes with detyrosi-nated tubulin, a marker of microtubule assembly. Hence, one possible avenue for future analyses will be to determine if *gdrd* functions in axoneme growth or stability. A major aspect of spermatogenesis that varies across *Drosophila* species is sperm tail length [69, 70]. While *gdrd* isn’t required for spermatid elongation, the gene may be required to generate or maintain long axonemes. Interestingly, we observed that the predicted structure of Gdrd protein in *Dmel* is largely unchanged from the predicted structure of Gdrd at the base of the *melanogaster* group, but differs from the predicted structure in outgroup species. One difference in sperm between the *melanogaster* group and its immediate outgroup, *obscura*, is that sperm are longer in the former set of species [69]. Thus, it is possible that Gdrd structural refinement is correlated with changes in overall sperm tail length.

The final level of functional analysis of *de novo* genes is a consideration of their evolutionary implications [31]. By definition, the ancestor in which Gdrd initially arose must have had the ability to produce sperm, so Gdrd was unlikely to be required for this process at its birth. In extant *Dmel*, however, the gene is completely essential for any sperm production. Furthermore, the gene is present in all species analysed except for *D. willistoni*, and its structure appears to be largely conserved since its origin. These data are consistent with two possibilities. First, Gdrd might have quickly evolved an essential function in late-stage spermatogenesis but became dispensable in the *D. willistoni* lineage because of lineage-specific changes to this process in the ancestor of this species. Less is known about spermatogenesis in this species, though ultrastructural studies of the process indicate that it is broadly similar to *D. melanogaster* [71]. Further mechanistic investigation of spermatogenesis in *D. willistoni* may help generate hypotheses about why Gdrd was lost specifically in this lineage, while it was retained in other divergent *Drosophila* species. Second, it is possible that in ancestors of the *melanogaster* group, Gdrd played a neutral or slightly beneficial role, consistent with its maintenance. Gdrd might have, at that point, fully integrated into the cellular interaction network, possibly via binding to other proteins, for example through formation of a coiled-coil. Gdrd may not, however, have immediately carried out a function essential for sperm production. Such a gain of function might have arisen later, possibly modulated by changes at Gdrd’s termini.

Our structural, functional and evolutionary analyses provide novel insights into the early evolution of a putative *de novo* evolved gene and highlight the subtle changes it underwent as it evolved toward its current, essential role in *Dmel* spermatogenesis. Our work may therefore serve as a blueprint for future investigations into the phenomenon of *de novo* gene emergence and the functionalization of the proteins they encode. Our results are consistent with and complementary to several large-scale studies [16, 72, 11, 14] that show that after their initial birth and gain of expression, many *de novo* proteins evolve slowly with only minor structural changes. Future studies may advance our understanding of how *de novo* genes evolve their functions by focusing on shorter evolutionary time scales, including population-level data, well-resolved structures, and a broader spectrum of functional conditions under which not-yet-adapted *de novo* proteins are accommodated by highly complex and well-established cellular networks. Such knowledge will improve our understanding of the evolution of proteins in general and may aid in devising new strategies for their design in the lab.

## Acknowledgements

We thank Susan Hawat (Department of Plant Biochemistry and Biotechnology, WWU Muenster) and Prof. Simone König (Core Unit Proteomics) for analyzing our proteins via mass measurements, Prof. Frank M. Boeckler (Molecular Design & Pharm. Biophysics, University of Tuebingen) and his group for providing us with their TEV-plasmid, Prof. Phillip Selenko (FMP Berlin, now Weizmann Institute, Israel) for measuring the NMR sample and helping with the analysis of the HSMC.

P.H.P. and G.D.F. were supported by NSF CAREER grant #1652013 to GDF. Fly stocks were obtained from the Bloomington Drosophila Stock Center (supported by NIH grant P40OD018537), and plasmids were provided by the Drosophila Genomics Resource Center (supported by NIH grant 2P40OD010949).

B.H. received funding from the EU under the Horizon 2020 Research and Innovation Framework Programm No. 722610.

## Author Contributions

A.L., B.H. and E.B-B. designed structural research; A.L. and B.H. performed structural predictions and ancestral reconstruction; A.L. performed cloning, expression, purification, CD, TSA and NMR measurements; T.S. helped with CD measurement and interpreting data; A.L., B.H., and A.M.D. and C.J.J. ran MD simulations and evaluated results; A.M.D. helped interpreting NMR data; P.H.P. and G.D.F. designed *Drosophila* experiments. P.H.P. performed genetic experiments and cytological analyses. All authors wrote and approved the final version of the manuscript.

## Material and Methods

### Online data availability

Scripts and additional data for this paper are available online at Zenodo DOI: 10.5281/zenodo.4291827

### Computational Methods

#### Structural prediction and homology detection

For prediction of protein disorder and secondary structure we used the programs s2D [73], as well as PSIPRED and Quick2D [74, 75] as implemented in the MPI bioinformatics toolkit [76]. *Ab initio* tertiary structures were predicted using the QUARK server (https://zhanglab.ccmb.med.umich.edu/QUARK) [77, 35]. The top five predicted models from QUARK were aligned using SALIGN, and RMSD values calculated using the res_cur command in PyMOL [78]. Kyte-Doolittle hydrophobicity was calculated with ExPASy’s ProtScale [79] using a window size of 19 residues. Aggregation propensity was predicted using TANGO [80] and solubility was predicted using [81]. Phase separation was predicted using PLAAC, taking background amino acid frequencies from the *Dmel* proteome interpolated at 50% with experimental *S. cerevisea* frequencies, and a minimum domain length of 40 aa [82]. To investigate the likelihood of our predicted structures for Gdrd representing diverged forms of existing homologs which have already been structurally solved, we took the top five predicted models from QUARK and searched for similar structures in the nr-PDB-90 database using 3D-BLAST with an E-value threshold of 1E^-15^ [40, 41]. For additional structural searches, we used the mTM-align align server with default settings (https://yanglab.nankai.edu.cn/mTM-align) [42]

#### Ancestral Sequence Reconstruction

Orthologs of Gdrd in other *Drosophila* species were gathered by three iterations of PSI-BLAST with an e-value threshold of 0.005 [83]. Sequences from *Dvir, Dmoj*, and *Dgri*, previously identified by Gubala et al. [30], were subsequently included. T-COFFEE v8.97 [84] was used to carry out structure-guided sequence alignment. A species tree was downloaded from timetree.org, and RAxML was used to carry out ancestral sequence and gap reconstruction [85]. Sequences were reconstructed under the PROTGAMMAJTT model, and gaps were reconstructed separately using maximum parsimony, following the methods described by Aadland et al. [86]. Most probable ancestral sequences were extracted for the relevant nodes.

#### MD simulation

MD simulations were performed using Gromacs 2018.1 [37, 38, 39] using the top predicted Gdrd structure from the QUARK webserver as an input structure. Structures were prepared following the standard procedure outlined in the Gromacs manual and tutorial. Prior to simulation, the structure was solvated in a cubic box of SPC/E water with 10 Å clearance and the electrostatic charge neutralized by addition of sodium atoms, followed by energy minimization and equilibration in Gromacs. Three 200 ns simulations were run in an NPT ensemble using a V-rescale modified Berendsen thermostat at a temperature of 300 K and a Parrinello-Rahman barostat at a pressure of 1 atm, periodic boundary conditions, and a particle mesh Ewald summation with a grid spacing of 1.6 Å and fourth order interpolation. Simulation trajectories were analyzed using Gromacs and the VMD package [87].

## Experimental Methods

A table of all primers used can be found in the supporting material (Table S1)

### *In vivo* tests of Gdrd

#### Fly stocks

Flies were raised at 25°C on standard media. Fly stocks: *w*^*1118*^, *dj-GFP*.*/CyO* (BL5417), *Df(3L)ED4543* (BL8073), and *Vas-Cas9* (BL51323) were obtained from the Bloomington *Drosophila* Stock Center. *ProtB-GFP* (*Mst35b-GFP*/ [88]) was used to construct the *Prot:GFP, +*/*+* and *Prot:GFP*; *gdrd*^*1*^/*TM3* lines used in this paper.

#### CRISPR-Cas9

5’ and 3’ flanking CRISPR-Cas9 gene editing target sites, GGTGGAACGGGTGGACG-GAATGG and CCAAACTTGCTTTCATTCGGTCC respectively, were identified using CRISPR Target Finder[89]. Guide RNAs were constructed by cloning annealed primers into pU6-3-gRNA vector (*Drosophila* Genomics Resource Center; Kate O’Connor-Giles). We then used the co-CRISPR technique as described in Ge et al. [90] to generate and screen for mutations at the *gdrd* locus (Rainbow Transgenics).

#### Fertility assay

Single virgin males were collected and aged for six days before they were mated individually to three Canton-S females. Both males and females were removed after three days of mating. Progeny number was determined by counting the number of pupal cases on the side of each vial ten days after setting the cross.

#### Gene Expression

mRNA transcripts of *gdrd* and *CG5048, RpL32* and were detected in RNA preps of *w*^*1118*^ and *gdrd* mutant flies as previously described [30]. The following primers pairs were used to detect each gene: *Gdrd* RT F/R; *CG5048* RT F/R; *RPL32*RT F/R.

#### Transgene

We used Gibson Assembly (NEB) to generate the HA-tagged Gdrd rescue construct. Putative upstream regulatory regions *gdrd* CDS and putative downstream regulatory regions were PCR amplified using Q5 High Fidelity Polymerase (NEB) and the Gdrd Rescue F1/R1 and Gdrd Rescue F3/R3 primer pairs (**Table S1**) respectively. The 3X HA tag was amplified using from pTWH (*Drosophila* Genomics Resource Center; T. Murphy) using Gdrd Rescue F2/R2 primers. PCR fragments were then assembled into a XbaI/AscI linearized w+attB plasmid (Sekelsky, Addgene plasmid 30326). Tagged rescue construct was then phiC31 integrated into the PBacy[+]-attP-9AVK00020 (BL24867) docking site (Rainbow Transgenics).

#### Immunostaining, phalloidin labeling, and microscopy

Testes were dissected in PBS, fixed for 20 minutes in 4% paraformaldehyde diluted in PBS, and subsequently permeabilized with PBX (PBS with 0.1% Triton-X). HA-tagged Gdrd was detected using rabbit anti-HA (Cell Signaling Technology, C29F4) diluted at 1:100 in PBX+ 5% Normal Goat Serum. Following overnight incubation in primary AB, samples were washed with PBX and then incubated with anti-rabbit Alexa 488 conjugated secondary antibody diluted 1:200 in PBX + 5% NGS. (Life Technologies, A11008). Secondary AB was washed away with PBX. Actin based structures were visualized by incubating fixed samples with TRITC-conjugated phalloidin (1:200; Molecular Probes) diluted in PBX for 2 hours. Nuclear DNA was visualized by incubating tissues with ToPro-3 Iodide (1 mM solution diluted to 1:1000; Invitrogen) for 15 minutes followed by PBS washes. Samples were mounted in Vectashield mounting medium (Vector Laboratories). Images were acquired using a Leica SP5 confocal microscope (CTR6000; Leica Microsystems) with its accompanying software using N PLAN 20.0 × 0.40 DRY, HCX PL APO CS 40.0 × 1.25 OIL UV, and HCX APO CS 63.0 × 1.40 OIL UV objectives (Leica Microsystems). Images were processed and analyzed using ImageJ Fiji [91]. Mann-Whitney U test and student two sample t-tests were used to determine p-values.

### Expression, purification, and structural analysis

#### Cloning of Gdrd

We used genomic DNA from the Canton S wild-type strain of *Dmel* for PCR to amplify the Gdrd sequence (flybase CG13477), primer see Table S1. The forward primer contains a BamHI and the reverse primer a HindIII cleavage site. As stop codon we used TAA. We digested the PCR product with both restriction enzymes (FastDigest, Thermo Scientific) for 3 h at 37°C. As vector we used the pHAT2 vector from the EMBL vector database, Heidelberg. This vector contains an N-terminal 6x Histag and the restrictions sites mentioned above. We used the same procedure for digestion of the vector (1 h 37°C) and purified the cleaved vector from agarose gel. After purification of both vector and insert with the purification kit from Zymo Research, we ligated both with an insert:vector ratio of 4:1 using ligase (Thermo Scientific, 1 h, 22°C). The ligation mix was purified again (Zymo Research) and 2 *µ*L of the purified reaction mix was added to 50*µ*L of chemical competent *E. coli* TOP10 cells. The cells were incubated for 30 min on ice, followed by a 90 sec heat-shock at 42°C. 500 *µ*L of LB-Media (5 g yeast extract, 6 g tryptone, 5 g NaCl) was added to the bacterial cell suspension and it was incubated for 1 h at 37°C. After incubation the cells were spread on an agar-plate containing 50 *µ*g/mL ampicillin (AMP) and incubated at 37°C over night. 8 clones were picked from the plate and investigated through a colony PCR to check for correct insert. Clones bearing the insert were incubated over night in 5 mL of LB + AMP at 37°C. The DNA was purified from the cells using the MiniPrep-Kit from Thermo Scientific and send for sequencing to Microsynth, Seqlab, Germany. Last, the correct DNA sequence was cloned into different BL21 strains (BL21(DE3), BL21 Star(DE3), T7 Express, and pLysS) with the protocol mentioned above for expression.

Additionally a TEV-cleavage site was cloned into the pHAT2-vector between the 6xHistag and target protein to remove the expression tag before NMR. Cloning and preparation of Gdrd was done as mentioned above.

#### Test-Expression and purification of Gdrd protein

To identify in which BL21 strain the protein gets expressed we first performed text-expressions. We inoculated 10 mL of LB+AMP from a glycerol stock of all four BL21 cells and let grow until solution got turbid (6-8 h, 37°C). We than aliquoted the solutions into 3×3 mL and incubated for 30 min at different temperatures (37°C, 28°C, and 20°C) before adding IPTG for a final volume of 0.5 mM and shaking over night.

500 *µ*L of each cell culture was centrifuged (15000 rpm, 2min). Pellets were re-suspended and lysed in 50*µ*L of BugBuster/Lysonase mix (former Sigma Aldrich, now Merck) through vortexing for 10 min. After centrifugation the supernatant was mixed with the same volume of SDS-loading buffer (standard). The pellet was resuspended in 5x diluted Bugbuster, centrifuged and resuspended in 50 *µ*L SDS-loading buffer. 10 *µ*L of each fraction was loaded on an SDS-gel (200V, 45min) and dyed using InstantBlue. Strain and temperature showing the best result were used for large expression and purification. For pHAT2-Gdrd this was BL21 star cells at 28°C and for pHAT2-TEV-Gdrd BL21 star at 20°C.

We also tried to use an MBP, Strep, Strep-FH8, and C-terminal 6xHis tag. None of these variants lead to soluble protein. Either the protein was not expressed at all all or packed directly into inclusion bodies, from which refolding was also not successful.

For an expression of a larger amount of Gdrd a pre-culture of 5 mL 2xYT (10 g yeast extract, 12 g tryptone, 5 g Nacl)+AMP was inoculated from glycerole stock and incubated at 37°C over night. This culture was added to 1 L of 2xYT+AMP and incubated at 37°C until an OD_600_ of 0.4 to 0.6 was reached. The culture was incubated for an additional hour at the appropriate temperature (20°C or 28°C, respectively) before IPTG was added to a final concentration of 0.5 mM and expression was done over night. Cells were harvested via centrifugation (6000 rpm, 15 min, 4°C) and re-suspended in buffer A (20 mM Phosphate-buffer, pH = 7.2, 150 mM NaCl, 15 mM Imidazole) and EDTAfree ProteaseInhibitor cocktail was added (Roche). Cells were lysed using ultrasound (20 s 60% burst, 1 min pause, 6 cycles) and centrifuged for 30 min at 12000 rpm at 4°C to separate from cell debris. The supernatant was filtered through a 0.45 *µ*m syringe filter and a sample was taken for SDS-gel (S). The cell-free extract was loaded onto an HiPrep Ni^2+^ column (GE Healthcare) using an ÄKTA start. The flow through was collected and a sample was taken for SDS-Gel (FT). The column was washed with 5 CV of buffer A (collected, sample W for SDS-Gel). The protein was eluted from the column using an 50 % gradient of buffer B (like buffer A, however, imidazole concentration was 250 mM). The eluted protein was fractionalized and analyzed using SDS-PAGE. The appropriate protein bands were combined and the protein was measured via mass to check if the protein is correct (see online material Zenodo DOI: 10.5281/zenodo.4291827).

pHAT2-TEV-Gdrd was purified the same way and the protein were pooled. TEV protease was added to the protein (final concentration 1 mg/mL) and cleavage took place over night at room temperature. After cleavage the buffer was exchanged with buffer C (20 mM Phos-phatebuffer, pH = 7.2, 150 mM NaCl) to remove imidazole using 3 kDa Amicon-centricons (Merck). The protein was then re-loaded onto the Ni-column to remove the His-tagged TEV protease. However, we encountered some stability issues during the cleavage process. While incubating the protein with TEV protease Gdrd tend to precipitate quite quickly after losing the His-tag. We tried the cleavage process under different conditions (4°C, 20°C, low salt, high salt, shorter cleavage time, different TEV concentrations, etc.). None of them helped to prevent the protein from precipitation. As consequence, we lost around 60 to 80 % of protein during this step.

Since the protein contains no cysteine residues, we purified Gdrd without using reducing conditions. We recovered soluble protein (8 mg/mL for tagged). Confirmation of the correct protein mass was determined by ESI-MS and trypsin maldi-tof.

#### CD measurements

Protein sample was transferred into buffer D (20 mM Phosphatebuffer, pH = 7.2, 150 mM NaF, chlorid free buffer) using Amicon centricons. CD spectrum were conducted on a Jasco J-815 spectrometer with a Jasco PTC-348WI Peltier type temperature control system (Jasco Corp, Hachioji, Japan) at constant nitrogen flow. Far-UV CD spectra were measured with 1 mm path length quartz cuvette. Gdrd was recorded at a concentration of 50 *µ*M. Spectrum was recorded from 190 nm to 260 nm with a resolution of 1 nm (50 nm/min). The final spectrum was corrected by subtracting corresponding baseline spectrum. To estimate the protein secondary structure from the measured CD spectra we used the K2D3 web server (http://cbdm-01.zdv.uni-mainz.de/andrade/k2d3/) [45].

#### NMR measurements

Overnight culture of 25 mL of pHAT2-Gdrd and pHAT2-TEV-Gdrd in BL21 star cells in 2xYT was centrifuged (6000 rpm, 5min) and re-suspended in 10 mL of 1 L of M9 medium (standard protocol) containing 1 mg of 15NH^4^Cl. Purification protocol was the same as mentioned above. The protein was measured on an Bruker-Biospin 600 MHz NMR by Phil Selenko, FMP Berlin, now Weizmann Institute, Israel. The NMR spectrum collected was processed using NMRViewJ (OneMoonScientific) and peak integration was performed using TopSpin v3.5 (Bruker) using a Lorentzian lineshape fit.

#### Thermal shift assay

The melting point of Gdrd was determined by a thermal shift assay (TSA). Either 50 *µ*M or 20 *µ*M purified Gdrd protein (in buffer C) was mixed with SYPRO orange 200x diluted in dimethyl sulfoxide (DMSO) rendering a final concentration of 5 % DMSO and 10X SYPRO orange. As blank buffer C was mixed with 200x SYPRO orange in DMSO. Denaturation curves were measured in 96-well plates in a Roche LightCycler 480 II using wavelengths of 465 and 580 nm for excitation and emission. A linear slope corrected sigmoid was fitted to the data and used to determine the melting temperature. For each Gdrd concentration, the measurements were performed six times (12 measurements in total) and the thermal transition temperature was calculated as the mean of all 12 measurements after blank subtraction (blank measure in triplicate) +/-1 standard deviation.

