## Supplementary Material for "Structural and functional characterization of a putative *de novo* gene in *Drosophila*"

### Supplemental material for "Structural and functional characterization of a putative *de novo* gene in *Drosophila*"

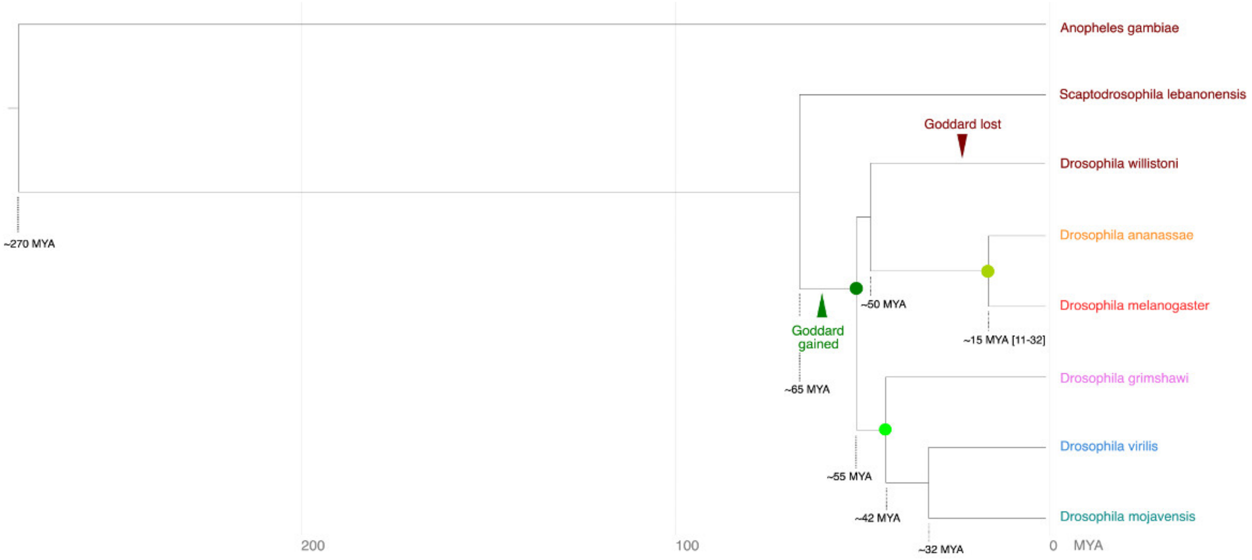


**Figure S1:** Phylogenetic tree showing the origination of the *gdrd* gene at the base of *Drosophila*. Gdrd appears to have emerged within an intron of the highly conserved omega gene, 50 million years ago. We previously ruled out the presence of the *gdrd* coding sequence in outgroup genomic regions syntenic to this intron [1]. Similarly, applying the same methodology, we find no evidence for an established *gdrd* gene in the recently released *Scaptodrosophila lebanonensis* genome. This suggests that *gdrd* is no older than 65 My. Reconstructed ancestral nodes are marked with green circles (colour matching main text Figure 6). Divergence times are best estimates from Obbard et al. (2012) and Russo et al. (2013) [2, 3].

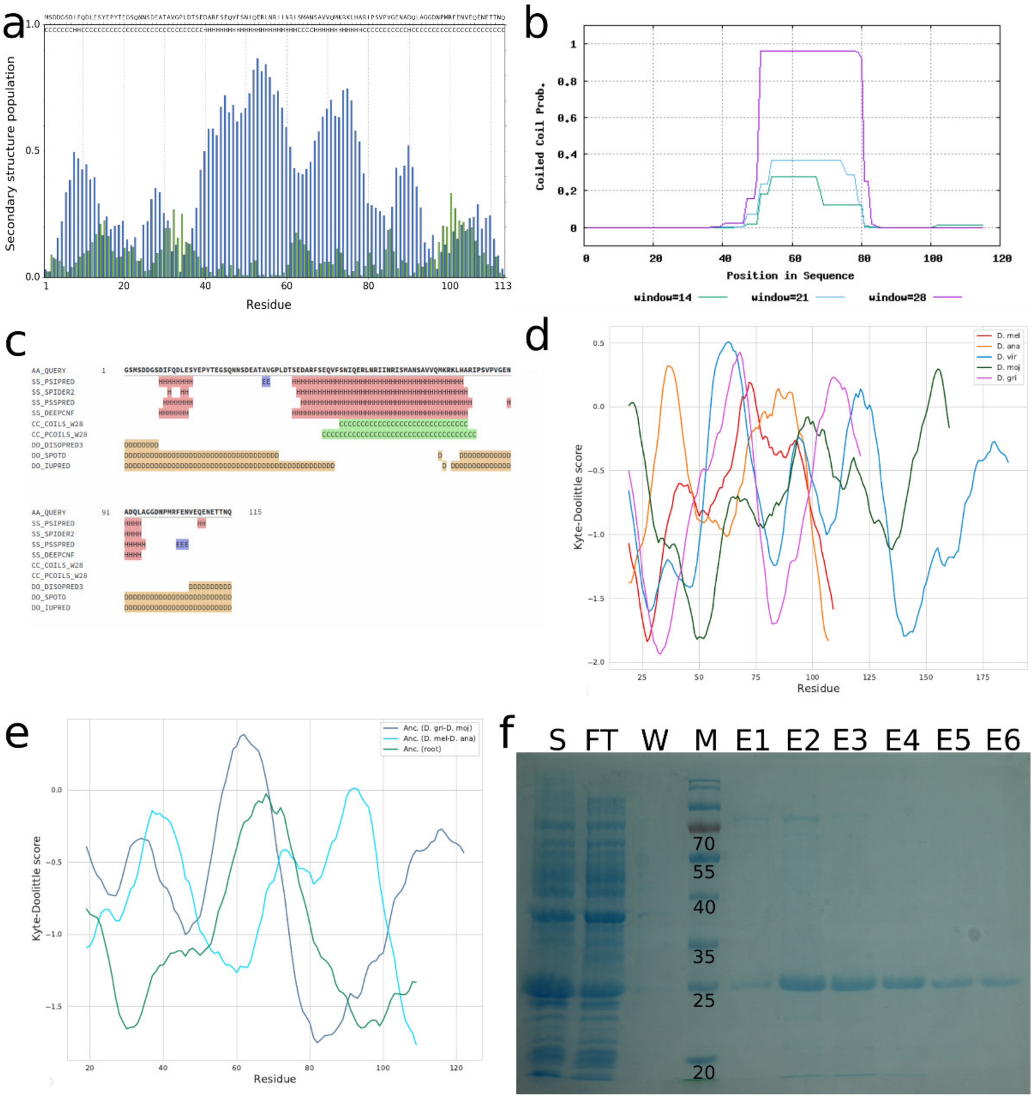


**Figure S2: Structural predictions for Gdrd** a) s2D[4], b) PCOILS[5], and c) Quick2D[6]. **Kyte-Doolittle plots illustrating the hydrophobicity** of d) Gdrd orthologs in all five *Drosophila* species and e) reconstructed sequences of Gdrd from three ancestral nodes. Regions of positive hydrophobicity are mainly isolated to the core of the protein in all sequences and correspond to helix predictions. f) **SDS-Gel of Gdrd purification**. Samples are (S) cell-free extract (before adding $Ni^{2+}$ beads); (FT) Flow through, (W) wash, (E1 to E6) elution steps with buffer B. Size of 6xHis-Gdrd is predicted to be 15kDa, however, always runs at 25 kDa in TGS-SDS-gels or BisTris gels. For mass detection (i) a band from elution fractions (for example like E2) was cut from SDS-Gel and analyzed via trypsin Maldi-Tof (Prof. König, Core Unit Proteomics, UKM Muenster) and (ii) a sample of combined elution fractions was taken for ESI-MS (Susan Hawat, Department of Plant Biochemistry and Biotechnology, WWU Muenster). Both measurements detected Gdrd protein (see mass data Zenodo DOI: 10.5281/zenodo.4291827).

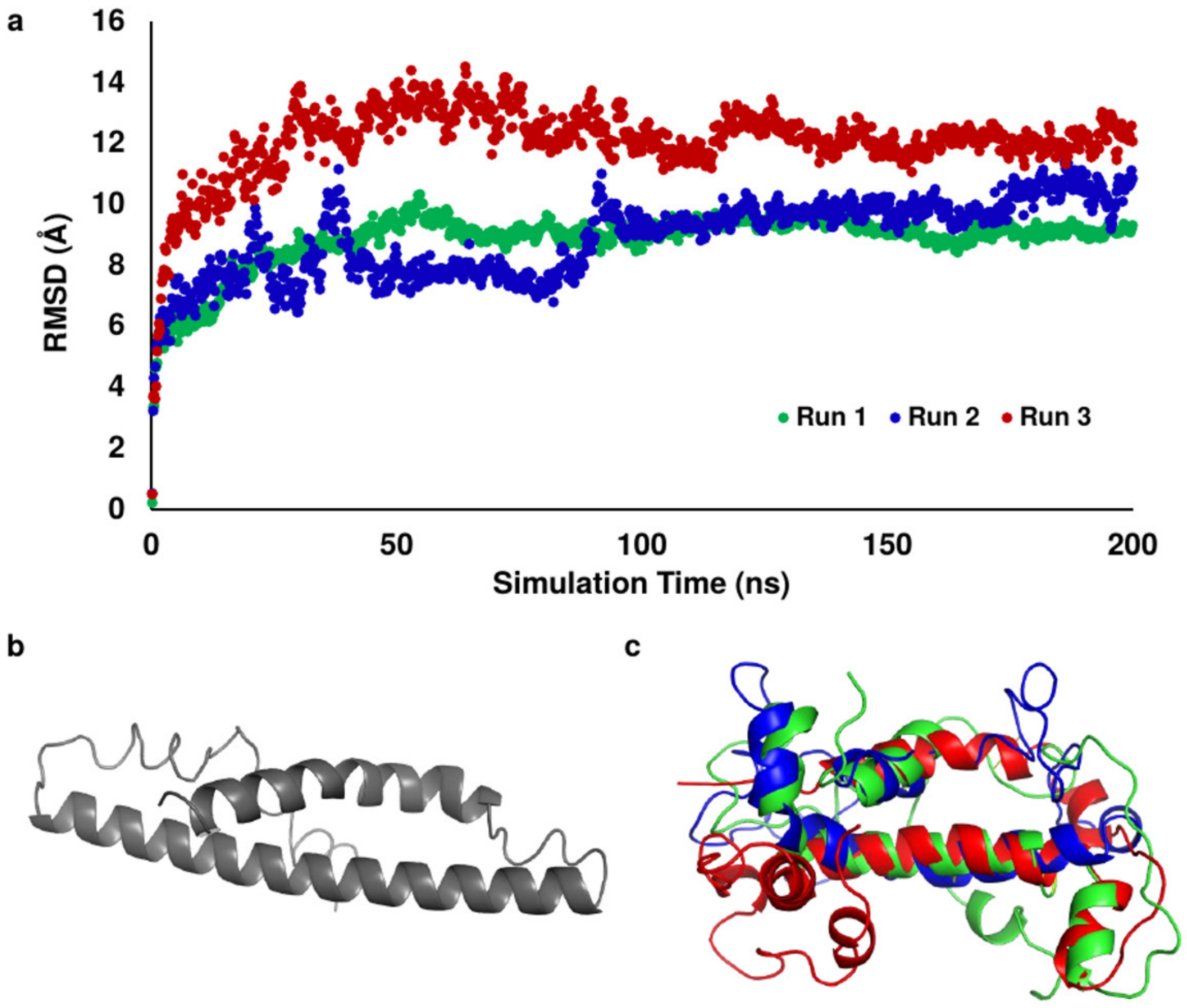


**Figure S3: MD simulation diversity.** a) Plot of simulation RMSD versus time shows a rapid divergence from the simulation input structure.[7, 8, 9] Following distortion of the starting loops and helices, all three trajectories reach relatively stable RMSD values within the first 100 ns of the simulation. b) Input structure obtained from the QUARK webserver[10]. c) Overlay of the endpoint structures from the three simulation replicates demonstrates the high variability observed in the protein C-terminus and loops. Positioning of the N-terminal helix and central helix however remains consistent.

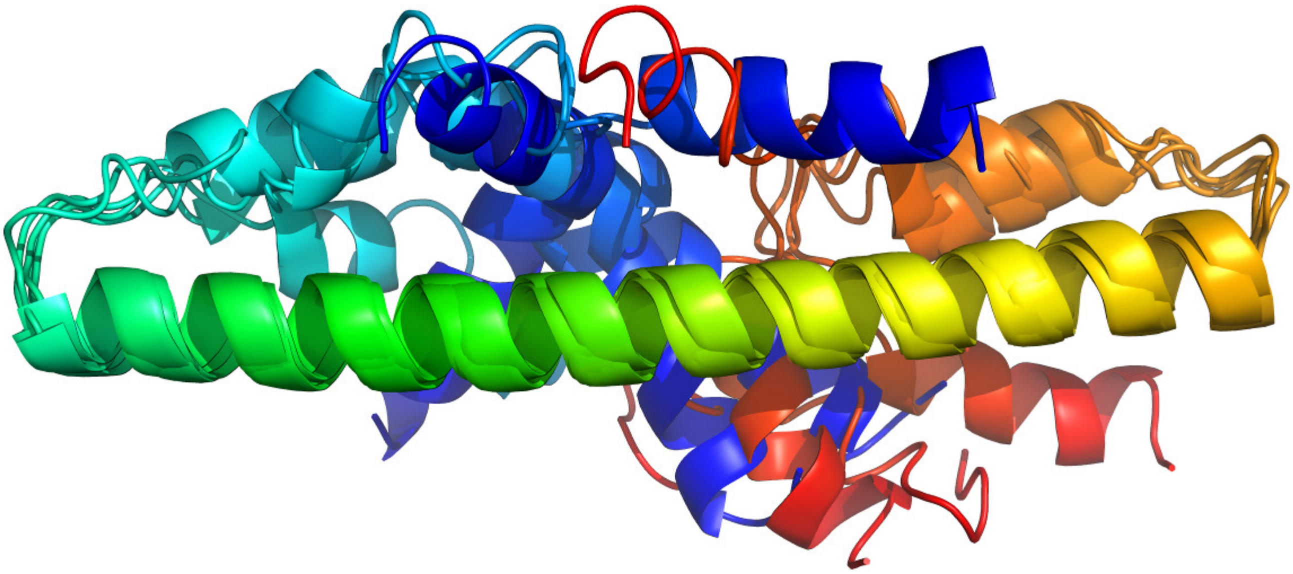

**Figure S4: Alignment of top 5 QUARK predicted structures of Gdrd.**[10] As expected, there are some differences at the termini of Gdrd, but the prediction of the core $\alpha$-helix is highly consistent (pairwise RMSD of 2.5 to 3.0 Å between residues 37-77). Pymol has been used for figure making.[11]

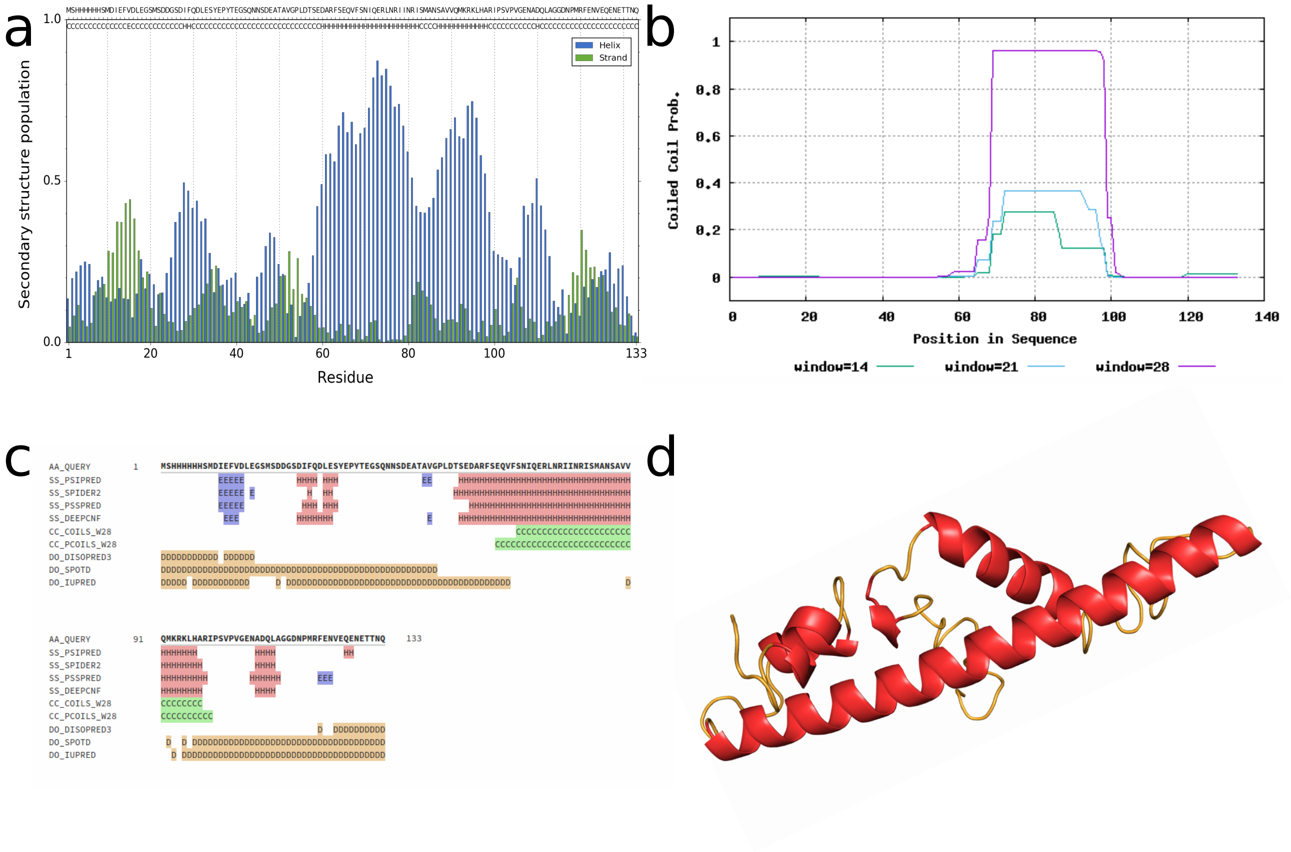


**Figure S5:** Structural predictions for the 6xHis-tagged *Gdrd* sequence using a) s2D[4], b) PCOILS[5], c) Quick2D[6], and d) QUARK (helix shown in red)[10]. Predictions are highly similar to the untagged version of Gdrd (Figure S1). The $\beta$-sheet region predicted by Quick2D (residues 15-18) is part of the 6x-His-tag.

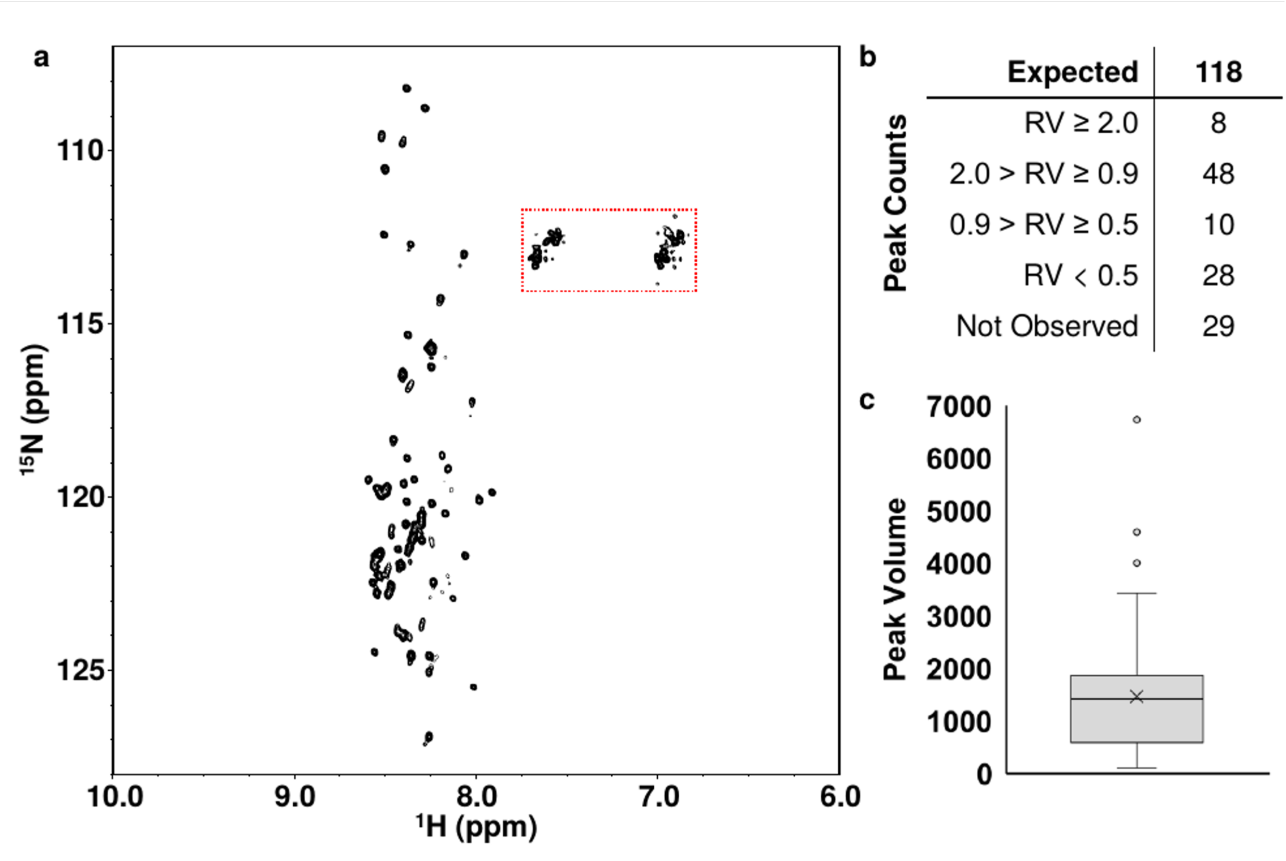


**Figure S6: NMR analysis of Gdrd suggests a partially-ordered helical structure.** a) $^{1}$H-$^{15}$N heteronuclear multiple quantum coherence (HMQC) spectrum of Goddard. The relatively poorly dispersed set of high-intensity peaks suggests that the ordered segments of the protein adopt a low-diversity secondary structure, which supports the modeled structure and MD simulations that suggest that these ordered regions are dominantly helical. Several additional peaks that are strongly broadened are also observed, suggesting that the remaining structure is highly flexible. Glutamine and asparagine side-chain peaks, boxed in red, were not included in further analyses. b) Count of peaks as a function of median-normalized relative volume (RV) demonstrates that roughly 40-50% of Goddard adopts an ordered structure (defined by the interval $2.0 \geq \text{RV} \geq 0.5$). c) Box plot of the peak volume distribution for observable peaks highlights the population of broadened, low-volume peaks that are indicative of conformational flexibility in the protein structure.

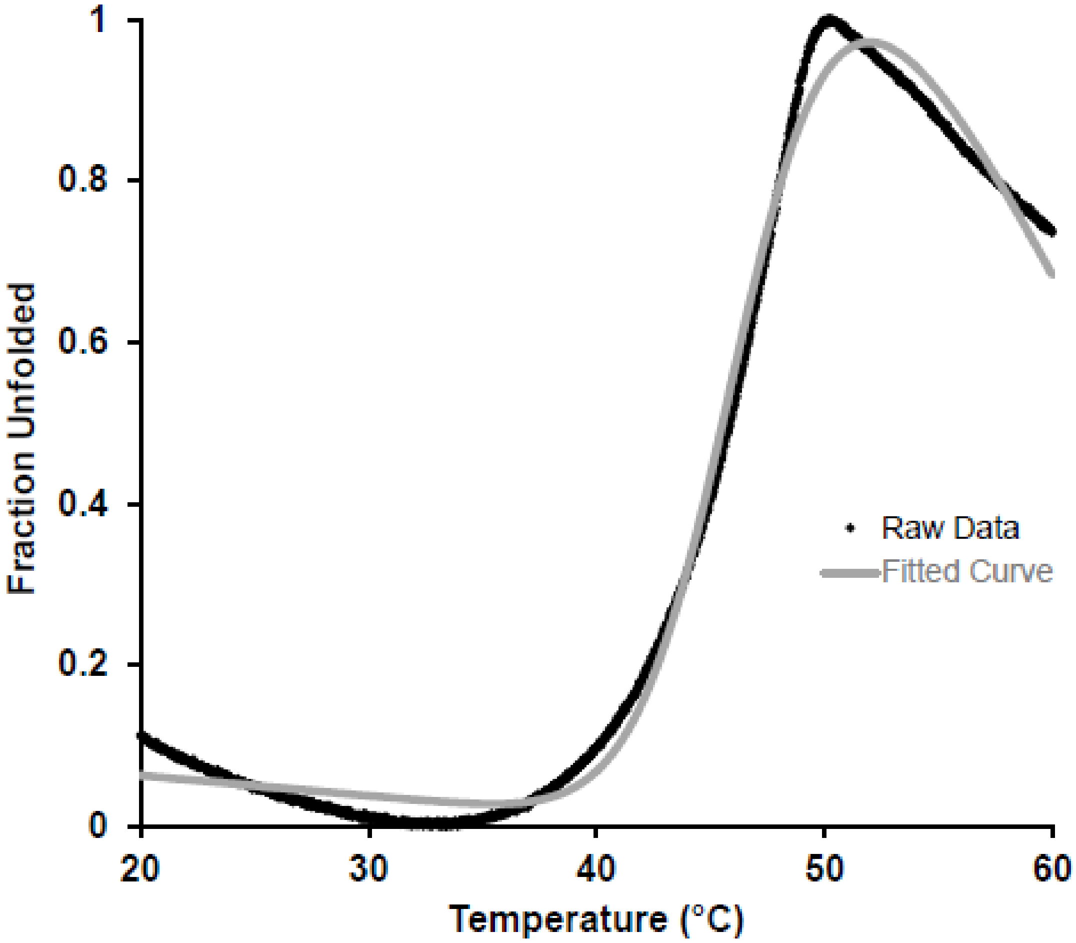


**Figure S7: Thermal unfolding assay of Gdrd**. Gdrd thermal unfolding data was obtained using a SYPRO orange based unfolding assay (see Materials Methods). Raw fluorescence data was converted to a fraction unfolded value under the assumption that the lowest fluorescence signal obtained corresponds to a fully folded protein and the highest fluorescence signal to a fully unfolded protein with all dye binding sites exposed. A linear slope corrected sigmoid was fit to the data and used to determine the melting temperature, 47.3°C +/- 0.9°C (average of n 12 =replicates +/- 1 standard deviation).

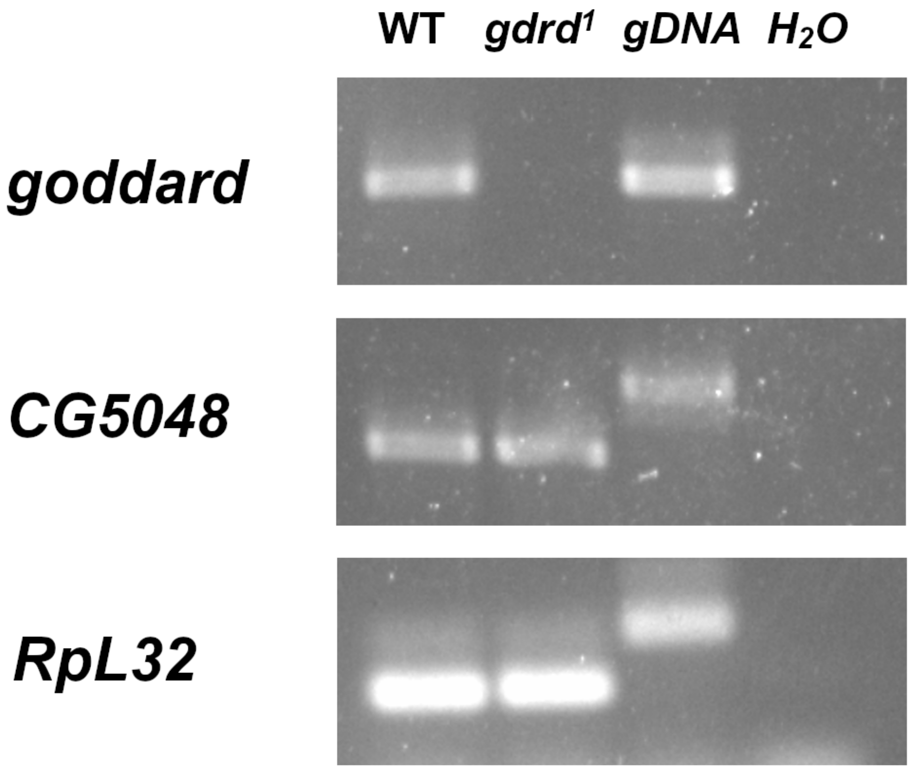


**Figure S8: *gdrd[1]* mutation specifically affects *gdrd* expression.** We performed RT-PCR analysis of *gdrd*, *CG5048*, and *RpL32* mRNA levels in wild type ($w^{1118}$) and *gdrd[1] flies*. *gdrd* expression is undetectable in *gdrd[1]* flies, while the expression of adjacent gene, *CG5048*, is unaffected in the *gdrd[1]* deletion mutant. Amplification using genomic DNA (gDNA) and water were performed as positive and negative controls respectively.

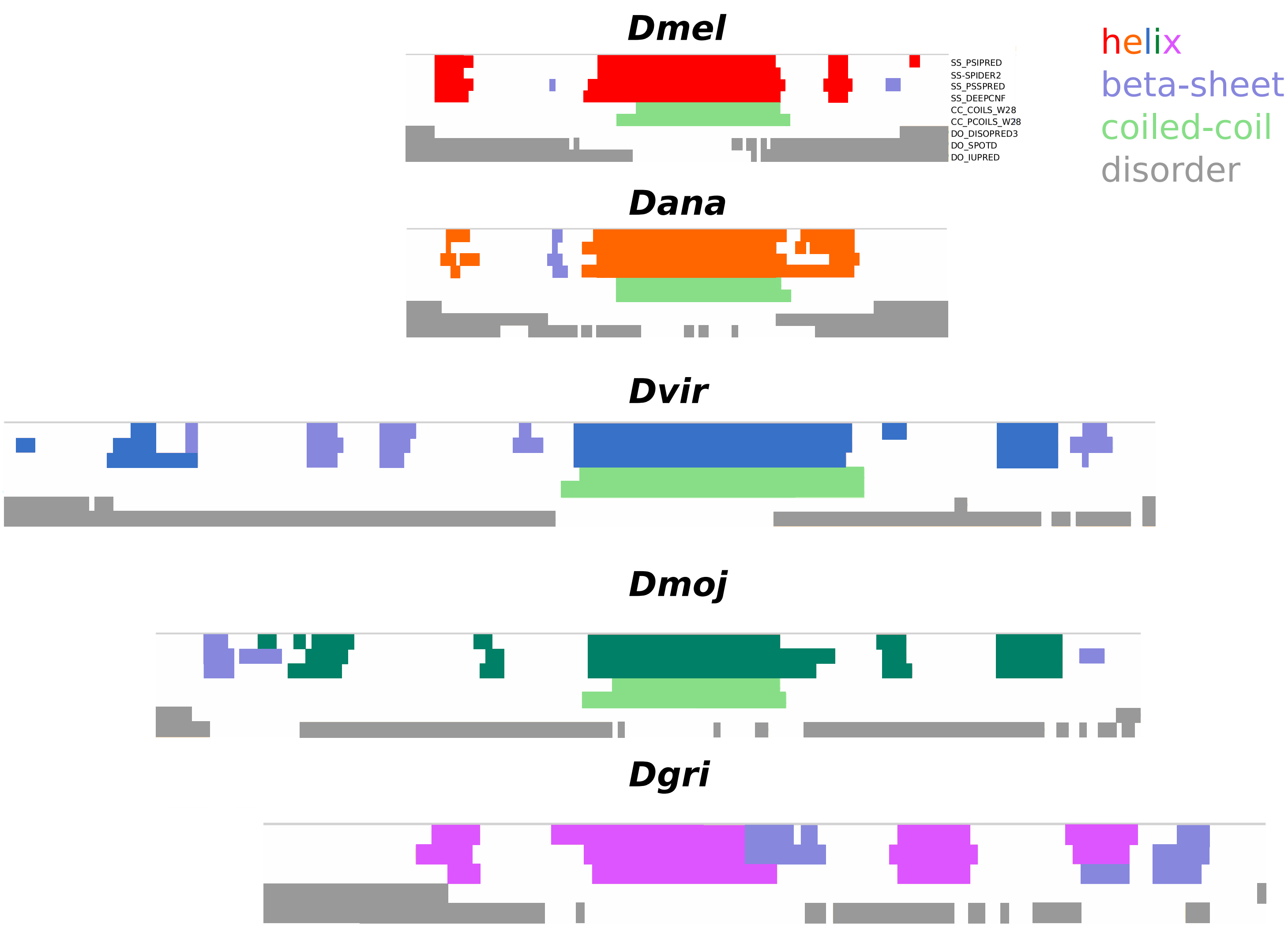


**Figure S9: Quick2D predictions of Gdrd orthologs from *Dmel*, *Dana*, *Dvir*, *Dmoj*, and *Dgr*.** Each block of color represents the linear amino acid sequence of Gdrd from different species. Helices are shown with a different colour in each species, $\beta$-sheets in lilac, disordered regions in grey [6].

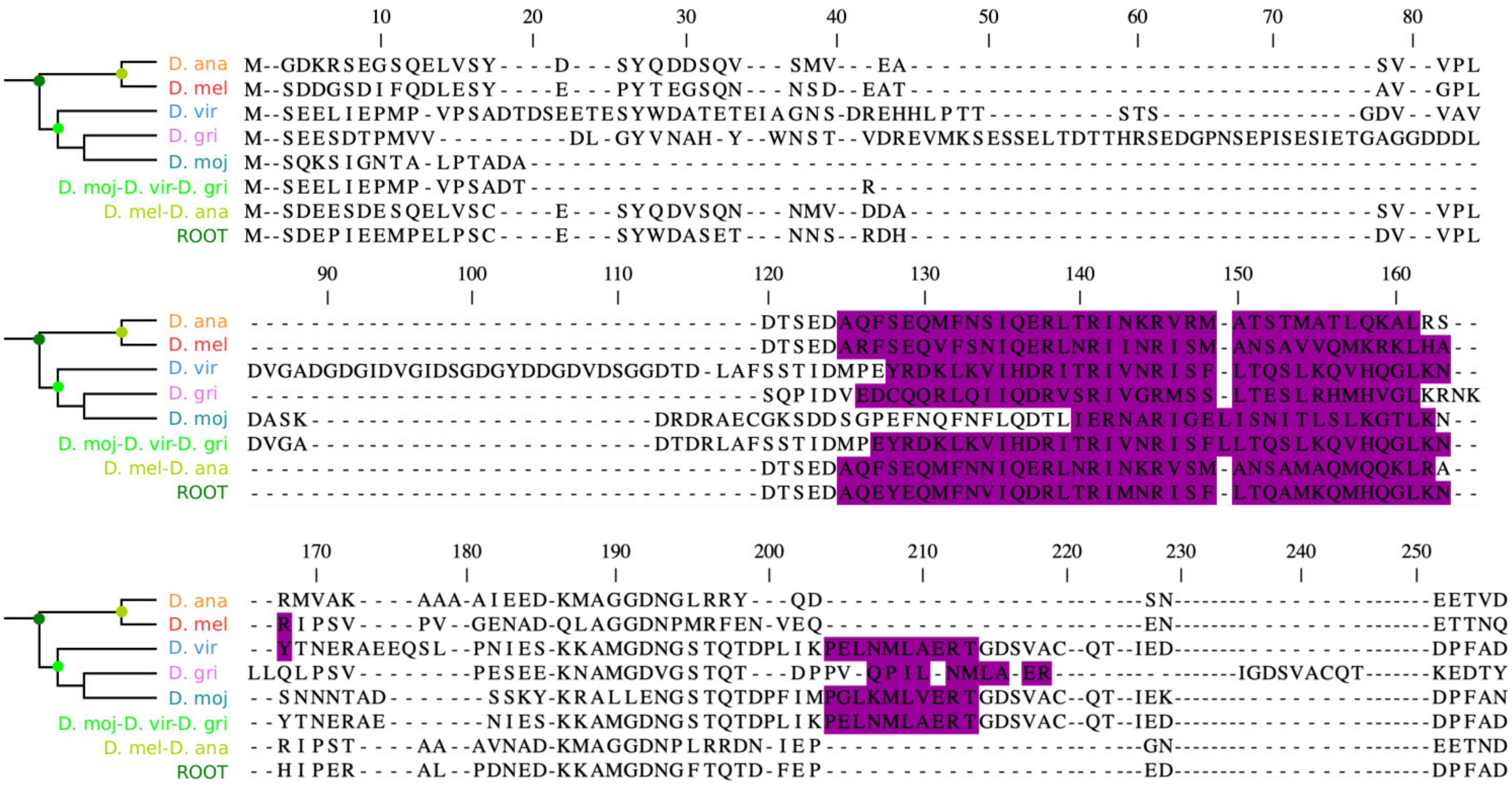


**Figure S10: Sequence alignment of Gdrd orthologs from five *Drosophila* species and three reconstructed ancestral sequences**. Secondary structure prediction using s2D[4] indicates highly conserved -helical character in extant and ancestral sequences (highlighted in purple).

| Primer name | Primer Sequence 5'– 3' | experimental use |
|---|---|---|
| Gdrd RT F | TCCAAGACCTTGAAAGCTACG | RT PCR |
| Gdrd RT R | TGGTTGGTGGTTTCGTTTTC | RT PCR |
| CG5048 RT F | GGAGTCAATGGTTGTCAATA | RT PCR |
| CG5048 RT R | ACATTTTGTCAGCAGTCATT | RT PCR |
| RPL32 RT F | CACCAGTCGGATCGATATGC | RT PCR |
| RPL32 RT R | CGATCCGTAACCGATGTTG | RT PCR |
| Gdrd Rescue F1 | GGCATGTCGACCTCGAGTACC CGGGAGCTCGAATTCTAGATA AGGAAAAGCGAAGCACAC | Tagged gdrd rescue (for amplification of upstream regulatory sequences and CDS) |
| Gdrd Rescue R1 | ATGGGTAAAAGATGCGGCCTC CACCGCGGTGGAGATCCATTT GGTTGGTGGTTTCGTTTT | Tagged gdrd rescue (for amplification of upstream regulatory sequences and CDS) |
| Gdrd Rescue F3 | TGATATAAGAACATTTTTATA TTTCTCATTTTCAAAAATGTATA AATTTATTGTATTTAT | Tagged gdrd rescue (for amplification of downstream regulatory sequences) |
| Gdrd Rescue R3 | ATTGCCGGCGATATCGGATCC ACCGGTGCCTAGGCGCGCCTG CAGAAGATGATTTAGAAA | Tagged gdrd rescue (for amplification of downstream regulatory sequences) |
| Gdrd Rescue F2 | ATGGATCTCCACCGCGGTGGA GGCCGCA | Tagged gdrd rescue (for amplification of HA tag) |
| Gdrd Rescue R2 | CATTTTTGAAAATGAGAAATAT AAAAATGTTCTTATATCACGTG GACCGGTGTCCGCCAT | Tagged gdrd rescue (for amplification of HA tag) |
| Gdrd F | GCGCGCGGATCCATGTCGGAC GACGGATCTGATATAT | For bacterial expression/purification of Gdrd (BamH1 site) |
| Gdrd R | CGCGCGAAGCTTTTATTGGTTG GTGGTTTCGTTTTCTT | For bacterial expression/purification of Gdrd (HindIII site) |

**Table S1:** Oligonucleotides used for molecular cloning and RT PCR.

[11] *The PyMOL molecular graphics system, version 1.2r3pre, schrödinger, LLC.*
